# Reducing PSY activity fine tunes threshold levels of a *cis*-carotene-derived signal that regulates the PIF3/HY5 module and plastid biogenesis

**DOI:** 10.1101/2023.06.29.546996

**Authors:** Xin Hou, Yagiz Alagoz, Ralf Welsch, Matthew D Mortimer, Barry J. Pogson, Christopher I. Cazzonelli

## Abstract

PHYTOENE SYNTHASE (PSY) is a rate-limiting enzyme catalysing the first committed step of carotenoid biosynthesis, and changes in PSY gene expression and/or protein activity alter carotenoid composition and plastid differentiation in plants. Here we identified four genetic variants of *PSY* (*psy*^−4^, *psy*^−90^, *psy*^−130^ and *psy*^−145^) using a forward genetics approach that rescued leaf virescence phenotypes displayed by the *Arabidopsis* CAROTENOID ISOMERASE (CRTISO) mutant *ccr2* (*carotenoid and chloroplast regulation 2*) when grown under a shorter photoperiod. The four non-lethal mutations affected alternative splicing, enzyme-substrate interactions, and PSY:ORANGE multi-enzyme complex binding, constituting the dynamic posttranscriptional fine-tuning of PSY levels and activity without changing localization to the stroma and protothylakoid membranes. *psy* genetic variants did not alter overall total xanthophyll or cis-carotene accumulation in *ccr2 yet* reduced specific acyclic linear *cis*-carotenes linked to the biosynthesis of a yet-to-be-identified apocarotenoid signal. *ccr2 psy* variants modulated the ratio of PHYTOCHROME-INTERACTING FACTOR 3/ELONGATED HYPOCOTYL 5 (PIF3/HY5), displayed a normal PLB formation in etioplasts, and chlorophyll accumulation during seedling photomorphogenesis. Thus, suppressing PSY activity and impairing PSY:ORANGE protein interactions reveals how threshold specific *cis*-carotene levels can be fine-tuned through holoenzyme-metabolon interactions to control plastid development.

**Highlights:** Manipulation of the PHYTOENE SYNTHASE catalytic activity in concert with its regulatory protein, ORANGE, reduces threshold levels of acyclic linear *cis*-carotenes that signal control over plastid biogenesis in dark and light grown Arabidopsis seedlings

## INTRODUCTION

Carotenoids are a diverse group of hydrophobic isoprenoid compounds synthesized by all photosynthetic and some non-photosynthetic organisms (Baranski and Cazzonelli, 2016). Carotenoid pigments provide colour and aroma to flowers, fruits and vegetables, attracting animals and insects for pollination and seed dispersal (Cazzonelli, 2011). Apart from very few insect species, animals are unable to biosynthesis carotenoids, and their ingestion by animals provides the substrate precursors needed to synthesise vitamin A (retinol), antioxidants, and derivative signalling metabolites that promote human health and immunity (Cazzonelli *et al*., 2010a). Intermediate substrates can be enriched in specific mutant tissues where they are reported to function as signals, yet the enzymatic steps in the pathway that can regulate their threshold levels remains unclear.

PHYTOENE SYNTHASE (PSY) catalyses the first committed step in the carotenoid biosynthetic pathway through the formation of 15 *cis*-phytoene (phytoene) from two geranylgeranyl diphosphate (GGPP) molecules (Zhou *et al*,. 2022). Phytoene is then converted to all-*trans* lycopene in four steps catalysed by two desaturases, PHYTOENE DESATURASE (PDS) and ζ-CAROTENE DESATURASE (ZDS), and two *cis*-*trans* isomerases, 15-*cis*-ζ-CAROTENE ISOMERASE (ZISO) and CAROTENOID ISOMERASE (CRTISO) (Alagoz *et al*., 2018). CRTISO and light limit flux through the lycopene branch point in the carotenoid pathway together with epsilon- and beta-lycopene cyclase enzymes that modulate α-carotene and β-carotene biosynthesis. A series of hydroxylation and/or epoxidation steps subsequently converts α-carotene to lutein and β-carotene to zeaxanthin, antheraxanthin, violaxanthin and to neoxanthin, collectively comprising the most abundant carotenoids found in photosynthetic leaves (Baranski and Cazzonelli, 2016) (Fig. S1). The loss-of-function of most enzymatic steps in the upper *cis*-carotene pathway do not induce plant lethality except for PSY, PDS, and ZDS, while the two isomersases rate-limit carotenoid accumulation in a light-dependent manner (Alagoz *et al*., 2018).

Carotenoid cleavage products, referred to as apocarotenoids, regulate plant development and modulate responses to biotic (e.g., insect herbivory) and abiotic (e.g., light and drought) stimuli (Hou *et al*., 2016; Moreno *et al*., 2021). Yet to be identified apocarotenoids can also be derived from acyclic linear *cis*-carotenes herein referred to as a *cis*-apocarotenoid signal (*cis*-ACS) (Alagoz *et al*., 2018; Anwar *et al*., 2021). Mutations in the Arabidopsis ZDS lead to the accumulation of phytofluene and ζ-carotene isomers that, via CCD4 cleavage, trigger a *cis*-ACS regulating *PHOTOSYNTHESIS ASSOCIATED NUCLEAR GENE* (*PhANG*) expression and chloroplast translation that leads to impaired chloroplast biogenesis, cellular differentiation and disruptions in leaf and flower meristem identity (Avendano-Vazquez *et al*., 2014; Escobar-Tovar *et al*,. 2021; McQuinn *et al*., 2023). A *cis*-ACS likely derived from di-*cis*-ζ-carotene, neurosporene, and/or prolycopene produced by the loss-of-function in *crtiso ziso* double mutants was shown to regulate *PhANG* expression and plastid development during skotomorphogenesis (e.g., formation of the prolamellar body; PLB) and photomorphogenesis (e.g., thylakoid structures within the chloroplast) (Cazzonelli *et al*., 2020). Neurosporene and/or prolycopene were also linked to the epistasis in tomato colour mutations that involved feedback regulation of *PSY1* expression by CRTISO during tomato fruit development (Kachanovsky *et al*., 2012). Which specific *cis*-carotene (absent or present) and rate-limiting enzymatic step(s) in the pathway can modulate their threshold levels and flux towards generating a *cis*-ACS remains unclear.

Alterations in *PSY* transcription and/or translation regulate plant carotenoid content (Alvarez *et al*,. 2016; Cao *et al*,. 2012; Fraser *et al*,. 2007; Maass *et al*., 2009; Welsch *et al*,. 2010). For example, the transcription of *PSY* is modulated by light signalling through multiple factors, including phytochrome-interacting factors (PIFs) and gibberellin (GA)-regulated DELLA proteins (Cheminant *et al*., 2011; Toledo-Ortiz *et al*., 2010). In Arabidopsis, the ORANGE protein (AtOR) post-transcriptionally regulates PSY levels through protein-protein interactions (Zhou *et al*., 2015). The localization of PSY has also been found to dramatically affect enzymatic activity (Welsch *et al*., 2000), and a single amino acid change can alter the localization of PSY and its protein activity (Shumskaya *et al*., 2012). There is a scarcity of information as to which amino acids of PSY can undergo substitution and alter activity.

After prolycopene biosynthesis and prior to the branch of the epsilon/beta-branch in the pathway, CRTISO regulates isomerisation together with light forming a unidirectional photoswitch that rate-limits *cis*-carotene levels (Alagoz *et al*., 2018; Nayak *et al*., 2022). CRTISO appears to be the key rate-limiting step inducing threshold levels of acyclic linear *cis*-carotenes that impair signalling of plastid biogenesis (Dhami *et al*,. 2022). During skotomorphogenesis, the etioplasts of seedling tissues develop prolamellar bodies (PLB), the characteristic paracrystalline membrane structure that defines etioplasts and accelerates photomorphogenesis upon illumination (Park *et al*., 2002; Rodriguez-Villalon *et al*., 2009).

We reported that a linear *cis-*carotene derived apocarotenoid signal (*cis*-ACS) generated by the Arabidopsis *crtiso mut a* (*nctcr2*; *carotenoid and chloroplast regulation 2*) that perturbed PLB formation in etioplasts as well as chlorophyll accumulation and chloroplast development that lead to leaf virescence in newly emerged leaf tissues (Cazzonelli *et al*., 2020; Park *et al*., 2002). The *cis-* ACS acted in parallel to the repressor of photomorphogenesis, DEETIOLATED 1 (DET1), to transcriptionally regulate the expression of *PHYTOCHROME INTERACTING FACTOR 3* (*PIF3*), *ELONGATED HYPOCOTYL 5* (*HY5*) and *PROTOCHLOROPHYLLIDE OXIDOREDUCTASE* (*POR*) during plastid development (Cazzonelli *et al*., 2020). We demonstrated that the *cis*-ACS accumulated in dark-grown tissues of *ccr2* or when plants were grown under a shorter photoperiod (Cazzonelli *et al*., 2020) (Fig. S1). A mechanism that can fine-tune *cis*-carotene levels when CRTISO activity is impaired and/or reduced and modulate threshold levels remains to be discovered.

We hypothesised that a step in the pathway before CRTISO should induce metabolic epistasis to regulate threshold levels of *cis*-carotene in a light-dependent manner. We performed a forward genetics screen to restore leaf greening to virescent *ccr2* foliar tissues *ccr2* grown under a short photoperiod. We mapped the casual mutations and confirmed the single nucleotide polymorphisms were responsible for the phenotype revision. We functionally characterised the roles of multiple PHYTOENE SYNTHASE variants controlling *cis*-carotene biosynthesis and plastid biogenesis in *ccr2* etiolated and de-etiolated seedlings. We investigated how the different lesions impact interactions between PSY and Arabidopsis ORANGE (AtOR), carotenoid accumulation, and by inference apo-carotenoid signalling that regulates plastid development during seedling transition from skotomorphogenesis to photomorphogenesis.

## Material and Methods

### Plant Materials

The *Arabidopsis thaliana* ecotype Columbia (Col-0) and ecotype Landsberg erecta (Ler-0) were used as wild-type controls in this study. The Arabidopsis *ccr2* mutant, carrying a G-to-A mutation in *crtiso* at the start of intron 9 that leads to miss-splicing (Park *et al*., 2002), was in Col-0 background. The *ccr2* mutant was subjected to forward genetics and a second site revertant screen through mutagenizing seeds in ethyl-methane sulfonate (EMS). EMS-treated seeds were sown in the soil, plants were grown and seeds were collected from pools of 5–10 M_1_ plants. Approximately 40,000 M_2_ seedlings from 30 pools of M1 seeds were screened for the emergence of green juvenile rosette leaves under a 10-h photoperiod. The revertant mutants of *ccr2* showing the emergence of green juvenile rosette leaves that were not virescent when grown under a 10 hr photoperiod are referred to as *rccr2* (revertant of *ccr2*)

### Plant growth conditions and treatments

For soil-grown plants, seeds were sown on seed raising mixture (Debco). Seeds were then stratified for 3 d at 4°C in the dark before transferring to an environmentally controlled growth chamber set to 21°C and illuminated by approximately 120 mmol.m^-2^.sec^-1^ of fluorescent lighting. Unless otherwise stated, plants were grown in a 16-h photoperiod. Photoperiod shift assays were performed by shifting three-week-old plants grown under a 16-h photoperiod to a 10-h photoperiod for one week, and newly emerged immature leaves were scored as displaying either a yellow leaf (YL) or green leaf (GL) phenotype which reflect either impaired or normal plastid development, respectively.

For media-grown seedlings, Murashige and Skoog (MS) media (Caisson Labs; MSP01) was used with half-strength of Gamborg’s vitamin solution 1000X (Sigma Aldrich) and 0.5% phytagel (Sigma Aldrich), and pH was adjusted to 5.8. Arabidopsis seeds were sterilized for 3 h in chlorine gas in a sealed container, followed by washing seeds once with 70% ethanol and three times with sterilized water. Seeds were sown onto MS media and stratified for 2 d at 4°C in the dark to synchronize germination.

Etiolation experiments were performed by incubating stratified seeds on MS media in the dark at 21°C for 7 d, after which cotyledons were harvested under a dim green LED light. For de-etiolation experiments, stratified Arabidopsis seeds were incubated in the dark at 21°C for 4 d, then exposed to constant light (80 mmol.m^-2^.sec^-1^) for 72 h at 21°C. Cotyledon tissues were harvested at 24-h intervals for chlorophyll quantification.

Seed derived calli (SDC) were generated as previously described (Mathur and Koncz, 1998). Briefly, 5 mg surface-sterilized seeds were plated onto a 90 mm petri dish containing 40 mL of SDC medium (4.33 g.L^-1^ MS basal salts (GibcoBRL), pH 5.8, 3% w/v sucrose, 0.1% v/v Gamborg B5 vitamins (Sigma-Aldrich), 0.5 mg.L^-1^ 2,4-D, 2 mg.L^-1^ indole-3-acetic acid (Sigma-Aldrich), 0.5 mg.L^-1^ 2-isopentenyladenine (Sigma-Aldrich), 0.5% w/v phytagel). Seeds were germinated for 5 d under 16-h photoperiod and incubated for 16 d in the dark before harvesting tissues for HPLC analysis. Callus treated with norflurazon (NFZ) were transferred onto the same medium containing 1 µmol.L^-1^ NFZ prior to etiolation.

### Pigment quantification

Total chlorophyll was quantified as described previously (Porra *et al*., 1989) with minor modifications as previously described (Cazzonelli *et al*,. 2020). Reverse phase HPLC (Agilent 1200 Series) was performed using either the GraceSmart-C_18_ (4 mm, 4.6 x 250 mm column; Alltech) or YMC-C_30_ (250 x 4.6 mm, S-5mm) column as described (Alagoz *et al*., 2020). The C_18_ column was used to quantify β-carotene, xanthophylls and generate *cis*-carotene chromatograms, while the C_30_ column was used for an improved *cis*-carotene separation and absolute quantification. Carotenoids were identified based on retention time relative to known standards and their absorbance spectra at 440 nm (β-carotene, xanthophylls, pro-neurosporene, tetra-*cis*-lycopene), 400 nm (ζ-carotenes), 340 nm (phytofluene) and 286 nm (phytoene). Absolute quantification of xanthophyll pigments and acyclic linear *cis*-carotenes in micrograms per gram fresh weight (gfw) was derived from peak area using molar extinction coefficient and molecular weight as previously described (Anwar *et al*., 2022). HPLC-based pigment extraction from shoot-derived callus (SDC) samples was performed as previously published (Welsch *et al*., 2008). To examine the phytoene levels in yeast alongside split ubiquitin and β-galactosidase activity assays, yeast cells were lysed by sonication in 100% acetone on ice. Lysates were centrifuged at 4°C, and the following extraction of phytoene and HPLC analysis were performed as per described for SDC (Welsch *et al*., 2008).

### Transmission electron microscopy

Cotyledons from 5-d old etiolated seedlings were harvested in dim-green safe light and fixed overnight in primary fixation buffer (2.5% Glutaraldehyde, and 4% paraformaldehyde in 0.1 mol.L^-1^ phosphate buffer pH 7.2) under vacuum, post-fixed in 1% osmium tetraoxide for 1 h, followed by an ethanol series: 50%, 70%, 80%, 90%, 95% and 3 × 100%. After dehydration, samples were incubated in epon araldite (resin):ethanol at 1:2, 1:1 and 2:1, then 3 times in 100% resin. Samples were then transferred to fresh resin and hardened under nitrogen stream at 60°C for 2 d, followed by sectioning of samples using Leica EM UC7 ultramicrotome (Leica Microsystems). Sections were placed on copper grids, stained with 5% uranyl acetate, washed thoroughly with distilled water, dried, and imaged with H7100FA transmission electron microscope (Hitachi) at 100 kV. For each of the dark-grown-seedling samples, prolamellar bodies were counted from 12 fields on 3 grids.

### DNA-seq library construction, Sequencing and Bioinformatics Identification of SNPs

Genomic DNA was extracted using the DNeasy Plant Mini Kit (QIAGEN). One microgram of genomic DNA was sheared using the M220 Focused-Ultrasonicator (Covaris) and DNA libraries were prepared using NEBNext® Ultra™ DNA Library Prep Kit (New England Biolabs) and size selected (∼320 bp) using AMPure XP Beads (Beckman Coulter). Libraries were then pooled and paired-end sequencing (NGS) was performed using the Illumina HiSEQ1500. The total number of reads ranged between 29-45 million representing approximately 22-35 times genome coverage. After sequencing, the raw reads were assessed for quality using the FastQC software (http://www.bioinformatics.babraham.ac.uk/projects/fastqc/), and subjected to trimming of illumina adapters and filtering of low quality reads with Adapter Removal programme (Lindgreen, 2012). The reads were mapped to the Arabidopsis (TAIR9) genome with BWA mapper (Li and Durbin, 2009). The resultant BWA alignment files were converted to sorted bam files using the samtools v0.1.18 package (Li *et al*., 2009) and was used as input for the subsequent SNP calling analyses.

The SNPs were called and analysed further on both the parent and mutant lines using NGM pipeline (Austin *et al*., 2011) and SHOREmap (Schneeberger *et al*., 2009). For the NGM pipeline, the SNPs were called using samtools (v0.1.16) as instructed and processed into ‘.emap’ files using a script provided on the NGM website. The .emap files were uploaded to the NGM web-portal to assess SNPs with associated discordant chastity values. To identify mutant specific SNPs, Parental line SNPs-filtered for EMS changes and homozygous SNPs were defined based on the discordant chastity metric. For SHOREmap, the SHORE software (Ossowski *et al*., 2008) was used to align the reads (implementing BWA) and called the SNPs (Hartwig *et al*., 2012). SHOREmap backcross was then implemented to calculate mutant allele frequencies and filter out parent SNPs and defined the EMS mutational changes. Where appropriate, custom scripts were used to identify mutant specific, EMS SNPs, to filter out parent SNPs and annotate the region of interest.

The localization of SNPs and InDels were based on the annotation of gene models provided by TAIR database (http://www.arabidopsis.org/). The polymorphisms in the gene region and other genome regions were annotated as genic and intergenic, respectively. The genic polymorphisms were classified as CDS (coding sequences), UTR (untranslated regions), introns and splice site junctions according to their localization. SNPs in the CDS were further separated into synonymous and non-synonymous amino substitution. The GO/PFAM annotation data were further used to functionally annotate each gene.

### Plasmid construction

pEARLEY::PSY-OE binary vectors were designed to overexpress wild type Arabidopsis *PSY* cDNA fragment which was driven by the constitutive CaMV35S promoter. The full-length cDNA coding region was chemically synthesised (Thermo Fisher Scientific) and cloned into the intermediate vector pDONR221. Next, using gateway homologous recombination, the cDNA fragment was cloned into pEarleyGate100 vector (Earley *et al*., 2006). Vector construction was confirmed by restriction digestion and Sanger sequencing.

To express recombinant PSY proteins in *E. coli*, 55 codons encoding a predicted chloroplast-targeting signal were deleted from the 5’-end of the *PSY* cDNA fragments. The cDNA fragments were then codon-optimised and cloned into pRSETA (ThermoFisher Scientific) through restriction sites *Xho* I and *Eco*R I that were attached respectively to the 5’- and 3’-end of the chemically synthesized sequences. To facilitate the purification of recombinant PSY without additional tags, sequence “GAAAATCTGTATTTTCAGGGT” that encodes TEV protease (Tobacco Etch Virus nuclear-inclusion-a endopeptidase) was inserted between *Xho* I site and start codon. Chemical synthesis and cloning were carried out by ThermoFisher Scientific, and the plasmids were verified by sequencing and restriction enzyme digestion. pRSET-PSY-WT, pRSET-PSY4, pRSET-PSY90 and pRSET-PSY130 were transformed to *E. coli* strain DH5α for propagation, followed by transforming BL21 (DE3) strain for the expression of recombinant PSY.

To transiently express recombinant PSY or psy variants in *Arabidopsis* etioplasts, full length coding regions of PSY, psy^−4^, psy^−90^, psy^−130^ and psy*-*N_181_P_270_ were chemically synthesized by Thermo Fisher Scientific and cloned into the intermediate vector pDONR221, followed by gateway cloning to pDEST15-CGFP vector (Gift from Professor James Whalen, La Trobe University) to construct pCGFP-AtPSY-WT, pCGFP-AtPSY4, pCGFP-AtPSY90, pCGFP-AtPSY130 and pCGFP-AtPSY-NP.

### Generation of transgenic plants

The *ccr2 psy* ^−4^*, ccr2 psy* ^−90^*, ccr2 psy* ^−130^ *and ccr2 psy* ^−145^ EMS-generated double mutant lines were transformed by dipping Arabidopsis flowers with Agrobacteria harbouring pEARLEY::PSY-OE binary vector to generate *ccr2 psy* ^−4^::PSY-OE, *ccr2 psy* ^−90^::PSY-OE, *ccr2 psy* ^−130^::PSY-OE and *ccr2 psy* ^−145^::PSY-OE transgenic lines, respectively (Clough and Bent, 1998). At least 10 independent transgenic lines were generated for each double mutant by spraying seedlings grown on soil with 50 mg.L^-1^ of glufosinate-ammonium salt (Basta herbicide).

### Expression, purification and identification of recombinant proteins

*E. coli* strain BL21 (DE3) cells harbouring pRSET-PSY-WT, pRSET-PSY4, pRSET-PSY90 or pRSET-PSY130 vector were inducted by adding 0.5 mmol.L^-1^ IPTG (Isopropyl β-D-1-thiogalactopyranoside, final concentration) and constant shaking at 28°C. Cells were harvested after 4-h induction by centrifugation. The pellets were washed with cold PBS and resuspended in buffer A (50 mmol.L^-1^ NaH_2_PO_4_, 20 mmol.L^-1^ imidazole, 1 mol.L^-1^ NaCl, pH 7.4), and lysed by lysozyme treatment and sonication on ice. After centrifugation, the supernatant containing the soluble fraction was loaded onto a Pierce 5-mL Ni-NTA column (Thermo Fisher Scientific) equilibrated with buffer A. The column was washed with buffer A containing low-concentration imidazole. Protein was then eluted using buffer A containing 200 mmol.L^-1^ imidazole, 10 μL of which was subjected to gel electrophoresis and western blot with anti-PSY polyclonal antibody to identify the recombinant PSY proteins.

Buffer exchange to TEV digestion buffer (50 mmol.L^-1^ Tris.HCl pH 8.0, 150 mmol.L^-1^ NaCl, 20 mmol.L^-1^ KCl, 2 mmol.L^-1^ β-mercaptoethanol) was carried out using a Pierce desalting spin column (Thermo Fisher Scientific). The column was equilibrated with TEV digestion buffer and purified recombinant protein was loaded. Protein samples were then collected by centrifuging the column and pooled after buffer exchange. Protein concentration was determined using Bradford reagent (Bio-Rad), followed by digestion using TEV at 4°C for 14 h. A Pierce 5-mL Ni-NTA column (Thermo Fisher Scientific) was equilibrated with TEV digestion buffer and TEV-cleaved recombinant PSY was loaded, followed by elution of the recombinant protein with TEV digestion buffer. The concentration of TEV-cleaved and purified recombinant PSY protein was determined again before *in vitro* enzymatic assays.

### In vitro activity assay of recombinant PSY

To synthesise GGPP, *E. coli* strain TOP10 cells harbouring artificial chromosome pAC-GGPPipi were cultured in chloramphenicol-containing LB medium at 28°C for 18 h. TOP 10 strain carrying pAC-PHYT was used as a positive control to verify the absorbance peak of phytoene in HPLC.

Cells were harvested by centrifugation and 1 g of cells (wet weight) were resuspended in 2 mL of enzyme assay buffer (100 mmol.L^-1^ Tris.HCl pH 7.6, 10 mmol.L^-1^ MgCl_2_, 2 mmol.L^-1^ MnCl_2_, 1mmol.L^-1^ 3,3’,3’’-phosphanetriyltripropanoic acid, 20% v/v glycerol and 0.08% v/v Tween 80). Cells were lysed by sonication on ice, followed by centrifugation. To the supernatant, 5 μg of purified recombinant PSY protein was added, followed by incubation at 20°C in the dark for 20 min. At a 4-min interval, aliquots were withdrawn, and the reaction was stopped by adding EDTA. Assays were extracted by adding carotenoid extraction buffer and partitioning phytoene into the ethyl acetate phase. The extractions were dried under nitrogen stream in the dark, resuspended in ethyl acetate and subjected to HPLC analysis.

### In vitro and in vivo activity assay of endogenous Arabidopsis PSY

The immunoprecipitation of endogenous PSY was performed using P-PER® Plant Protein Extraction Kit and Pierce Co-Immunoprecipitation kit (both from Thermo Fisher Scientific) following manufacturer’s instructions and is summarised below.

For total protein extraction, 160 mg leaf tissue from a 25-d old Arabidopsis plant grown under 16-h photoperiod was homogenised in P-PER® Working Solution containing Halt™ Protease Inhibitor Cocktail (Thermo Fisher Scientific). Following centrifugation, the lower aqueous layer containing total protein was recovered for immunoprecipitation. Twenty micrograms of anti-PSY antibody was coupled to AminoLink Plus coupling resin and the total protein extracted from Arabidopsis leaf tissue was incubated with antibody-coupled resin overnight at 4°C. The resin was then washed and proteins were eluted with elution buffer, followed by neutralisation of the eluent by adding 1 mol.L^-1^ Tris (pH 9.5). Five micrograms of endogenous PSY co-immunoprecipitate (with binding proteins) was subjected to enzymatic activity assay. Immunoprecipitation with normal rabbit IgG (Agrisera Antibodies AS101545) was included as a negative control for the specificity of the anti-PSY antibody.

*In vivo* PSY activity was examined by measuring phytoene levels in norflurazon-treated Arabidopsis SDC. Generation of SDC and HPLC analysis was carried out as described in this study.

### Split Ubiquitin Protein-Protein Interaction Assay

The *PSY* cDNA sequences were truncated, and the 55 codons were deleted from the 5’-end. The cDNA sequences of Arabidopsis *OR* and *GGPS11* genes were obtained as previously described (Zhou *et al*., 2015). The truncated cDNA sequences of *PSY*, *OR* and *GGPPS11* were cloned to make Cub- or Nub-fusion constructs which were then transformed into yeast strain THY.AP5 (Cub) or THY.AP4 (Nub). Yeast strains carrying Cub fusions were mated with strains carrying Cub fusions, and the resulting diploid cells were grown on synthetic complete medium lacking leucine and tryptophan (-LW). Interaction growth tests were performed with overnight cultures spotted on fully selective medium (-LWAH), in a series of 1:10 dilutions with a starting OD_600_ of 2.0 after growing for about 2 d at 29°C. To reduce background activation of reporter genes and visualize different interaction strengths, methionine was added to the medium suppressing the expression of Cub-fusion proteins (+Met; 150 μmol.L^-1^ and 1 mmol.L^-1^). Control combinations with empty Cub- or Nub-expressing vectors were included in the experiments.

For β-galactosidase activity assays, yeast cells were pelleted from 250 μL overnight culture in complete medium and resuspended in 650 μL assay buffer (100 mmol.L^-1^ HEPES.KOH pH 7.0, 150 mmol.L^-1^ NaCl, 2 mmol.L^-1^ MgCl_2_, and 1% w/v BSA), followed by adding 50 μL chloroform and 50 μL 0.1% (w/v) SDS and vortex. Enzymatic reactions were started by adding 125 μL of 4 mg.mL^-1^ *ortho*-nitrophenyl-β-galactoside in the assay buffer. Following incubation at 30°C, until the mixture turned visibly yellow, reactions were stopped by adding 1 mol.L^-1^ Na_2_CO_3_. Reactions were then centrifuged, and OD_420_ was measured to determine the concentration of *ortho*-nitrophenol (oNP) in the supernatant. The Β-galactosidase activity was calculated as nmol oNP.min^-1^.OD_600_ ^-1^, using the molar extinction coefficient:oNP = 3300 g.L^-1^.mol^-1^. All β-galactosidase activity assays were performed in triplicate.

### Protoplast isolation and transient expression of recombinant PSY protein

To isolate etiolated Arabidopsis protoplasts, 200 mg of etiolated cotyledons were harvested under green safe light and treated in a 9-mm Petri dish with an enzyme solution (0.5% cellulase, 0.05% pectinase, 600 mmol.L^-1^ mannitol, 10 mmol.L^-1^ CaCl_2_ and 20 mmol.L^-1^ MES pH 5.6) at 21°C for 6 h with gentle shaking, followed by shaking vigorously for 3 min. The protoplasts were collected through a 60-μm nylon filter and washed 3 times with 20 mL cold washing buffer (600 mmol.L^-1^ mannitol, 10 mmol.L^-1^ CaCl_2_ and 20 mmol.L^-1^ MES pH 5.6), by centrifugation at 120× g for 5 min and resuspending the pellet. The number of protoplasts per mL was determined using a haemocytometer. In 150 µL of 600 mmol.L^-1^ mannitol containing 10 mmol.L^-1^ CaCl_2_, 20 µg of PSY-CGFP plasmid was added to 10^6^ protoplasts and mixed gently for 5 s. Five hundred microliters of 40% polyethylene glycol 6000 solution (in 500 mmol.L^-1^ mannitol and 100 mmol.L^-1^ Ca(NO_3_)_2_) was added and the contents were mixed gently for 15 s, followed by dilution with 4.5 mL mannitol/MES solution (500 mmol.L^-1^ mannitol, 15 mmol.L^-1^ MgCl_2_ and 0.1% MES pH 5.6) and incubation at 21°C for 20 min. The protoplasts were pelleted by centrifugation at 120× g and washed with 600 mmol.L^-1^ mannitol and 10 mmol.L^-1^ CaCl_2_, followed by incubation for 16 h at 25°C in the dark. The protoplasts were subjected to western blot using isolated stromal and membrane fractions.

### Isolation of stromal and membrane fractions from etioplasts

About 5× 10^6^ etiolated protoplasts transiently expressing recombinant PSY were centrifuged at room temperature at 250× g, and then at 4°C at 2,000× g. The pellet was resuspended in 10 mL isolation buffer and layered on 40% (w/v) Percoll® (Sigma-Aldrich) followed by centrifugation at 4°C. The Percoll® layer was removed, and the etioplast pellet was washed twice with isolation buffer. Etioplasts were resuspended in 600 µL hypotonic buffer (10 mmol.L^-1^ Tris.HCl pH 7.0 and 4 mmol.L^-1^ MgCl_2_) on ice for lysis. PLBs were then pelleted through centrifuging the lysate at 3,000× g at 4°C, and the supernatant was collected as the stromal fraction 10 µL of which was used for western blot. The pellet was resuspended in 5 mL hypotonic buffer and subjected to an ultrasonic bath treatment on ice. The suspension was layered on a sucrose step gradient (15 mL of 1 mol.L^-1^ sucrose under 10 mL of 600 mmol.L^-1^ sucrose in hypotonic buffer) and centrifuged at 4°C for 3 h at 75,000× g. Protothylakoids accumulating at the interface of the gradient were collected, diluted with an equal volume of hypotonic buffer and concentrated by ultracentrifugation at 125,000× g for 1 h at 4°C; the protothylakoids which pelleted below the gradient were harvested and combined with the fraction collected from the interface. The combined protothylakoid fractions were resuspended in 600 µL hypotonic buffer and 10 µL of each sample was used for western blot.

### Protein extraction and quantification

Fifty to one hundred milligrams of etiolated Arabidopsis seedlings (7-d old) were harvested under dim-green safe light ground to fine powder, or around 100 mg green leaf tissues of 25-d old plants were harvested under normal light. Total protein was extracted using a TCA-acetone method as previously described (Mechin *et al*,. 2007). The concentration of protein was measured using Bradford reagent (Bio-Rad) and adjusted to 2 µg.µL^-1^. A serial dilution was used to determine western blot sensitivity for each antibody and determine the optimal concentration for quantification. To examine recombinant PSY protein from *E. coli* cell lysates, endogenous PSY from stromal or membrane fractions of Arabidopsis etioplasts, 10 µL of sample was used for gel electrophoresis; to examine endogenous PSY or OR protein levels in Arabidopsis leaf tissues, 10 µg total protein was used; to detect OR protein from co-immunoprecipitation, 10 µL of PSY co-immunoprecipitate was used; to examine PIF3 or HY5 protein levels, 5 µg total protein was used. Proteins run on a gel were transferred to a PVDF membrane (Bio-Rad). Membranes were blocked and then incubated with anti-PSY antibody (1:1000, Agrisera Antibodies AS163991), anti-OR antibody (1:1000, gift from Prof. Li Li, College of Agriculture and Life Sciences, Cornell University), anti-HY5 antibody (Agrisera Antibodies AS121867, 1:1000) or anti-PIF3 antibody (Agrisera Antibodies AS163954, 1:2000) for 2 h at room temperature. Membranes were then washed and incubated with HRP-conjugated Goat anti-Rabbit IgG (1:5000, Agrisera Antibodies AS09602) at room temperature for 90 min, or for PIF3 with HRP-conjugated Rabbit anti-Goat IgG (Agrisera Antibodies AS09605, 1:5000) for 90 min. Membranes were re-probed using anti-Actin poly-clonal antibody (Agrisera Antibodies AS132640, 1:3000) and HRP-conjugated Goat anti-Rabbit IgG (Agrisera Antibodies AS09602, 1:5000) for internal protein normalisation.

### Enzyme-linked immunosorbent assay (ELISA)

Clear Flat-Bottom Immuno 96-well plates (Thermo Fisher Scientific) were coated with Arabidopsis total protein (20 µg.mL^-1^) diluted in carbonate buffer (pH 9.6) at 4°C overnight, followed by washing the plates with PBST (pH 7.4). Plates were then blocked for 1 h at room temperature and washed with PBST. Plates were incubated with anti-PSY (1 µg.mL^-1^, Agrisera Antibodies AS163991) or anti-OR (1 µg.mL^-1^, gift from Prof. Li Li), washed and incubated with HRP-conjugated Goat anti-Rabbit IgG (0.4 µg.mL^-1^, Agrisera Antibodies AS09602). All antibodies were diluted in PBS-TY buffer PBS-TY buffer (PBS with 0.05% v/v Triton X-100 and 1% w/v yeast extract, pH7.4) and all incubations were performed at 37°C for 1 h. After antibody incubations and washing the plates, substrate solution containing 0.5 mg.mL^-1^ O-phenylenediamine dihydrochloride (Sigma-Aldrich) in phosphate-citrate buffer (pH 5.0) and 0.02% (v/v) H_2_O_2_ was added and incubated for 5 min at room temperature. The reactions were stopped with 2mol.L^-1^ H_2_SO_4_, and the absorbance at 492 nm was measured on a TECAN M1000PRO plate reader (Tecan Group). Solubilisation buffer diluted in carbonate buffer was used as blank control, and a serial dilution of purified recombinant PSY protein in carbonate buffer was used as a positive control. All ELISAs were done in triplicates.

### Quantitative RT-PCR (qRT-PCR)

The total RNA was extracted using Spectrum™ Plant Total RNA kit (Sigma-Aldrich) as per the manufacturer’s protocol. First strand cDNA synthesis was performed using 1 µg total RNA, Oligo dT18 primer and Transcriptor First Strand cDNA synthesis kit (Roche) as per the manufacturer’s instructions. The qRT-PCR was performed using 2 µL of primer mix (2 µM for each primer), 1 µL 1/15 diluted cDNA template, 5 µL LightCycler 480 SYBR Green I Master mix and 2 µL sterile milli-Q water. For each sample, three technical replicates for each of the three biological replicates were tested. The relative gene expression levels were calculated by using relative quantification (Target Eff Ct(Wt-target)/Reference Eff Ct(Wt-target)) and fit point analysis (Pfaffl, 2001). Protein Phosphatase 2A (PP2A; At1g13320) gene was used as reference gene for normalisation (Czechowski *et al*., 2005). *PP2A* has been validated against Cyclophilin (At2g29960) and TIP41 (At4g34270) as secondary reference genes for different Arabidopsis tissues (Cazzonelli *et al*., 2014; Cazzonelli *et al*,. 2010b). DNA plasmids harboring wild type transcripts or splice variant cDNA was used as a reference to generate a standard curve and optimise the qRT-PCR conditions. All primer sequences are listed in Table S1.

### Protein Modelling

The Arabidopsis PSY structural models were built using the Colabfold (Mirdita *et al*., 2022) implementation of Alphafold2 (Jumper *et al*., 2021) and ESMFold (Lin *et al*., 2023). The ColabFold structure was inferred from the Arabidopsis PSY amino acid sequence with the signal peptide removed, using the ‘no template’ method, and the predicted structure was relaxed using amber force fields. Other settings were as the default. Five structures were predicted and ranked according to the predicted local distance difference test (pLDDT) (Mariani *et al*., 2011) confidence measure with the highest ranked model selected with a score of 88. The ESMFold model was made using one copy and three recycles with model confidence pLDDT score of 87. While the ESMFold model had higher pLLDT scores for the N- and C-termini and the N-terminal variable loop region, the pLDDT scores for the conserved core of the enzyme were marginally lower (confidence data is included with both structural models in the supplementary materials). As such the ColabFold model was chosen, and the fold was validated by comparing the model to the crystal structure of Enteroxoxxus hirae dehydrosqualene synthase complexed with three Mg2+ ions and the Farnesyl thiopyrophosphate – FPS ligand (PDB accession: 5IYS). Structural alignments were made using PyMOL (Schrödinger, 2015) and alignments of the ColabFold model with the ESMFold model and 5IYS crystal structures are included in the supplementary materials. Figures were made using PyMOL.

## RESULTS

### Mutations in PSY reduce cis-carotene levels and restore plastid development in ccr2

Our forward genetics approach uncovered uncharacterised second-site mutations (referred to as revertant *ccr2*; *rccr2* ^−4^, *rccr2*^−90^, *rccr2*^−130^ and *rccr2*^−145^) capable of restoring leaf greening to *ccr2* plants that would otherwise show virescence reflected by a yellow leaf (YL) phenotype when grown under a short (10 h) photoperiod (Fig. 1A, Fig. S2A). All *rccr2* lines showed enhanced chlorophyll levels in young emerging leaves (∼2-fold compared to *ccr2*) and in cotyledons following de-etiolation matching WT levels (Fig. 1B-C) and yet displayed reduced foliar lutein levels like *ccr2* (see Figure 3B in Cazzonelli et al., 2020). In dark-grown etiolated seedlings, the four *rccr2* lines displayed a PLB structure similar to, or partially restored, when compared to WT (Fig. 1D-E). Therefore, restoration of PLB formation in etiolated *rccr2* tissues leads to normal chlorophyll accumulation in de-etiolated seedlings and foliar tissues from plants grown under a shorter photoperiod.

**Fig. 1:**
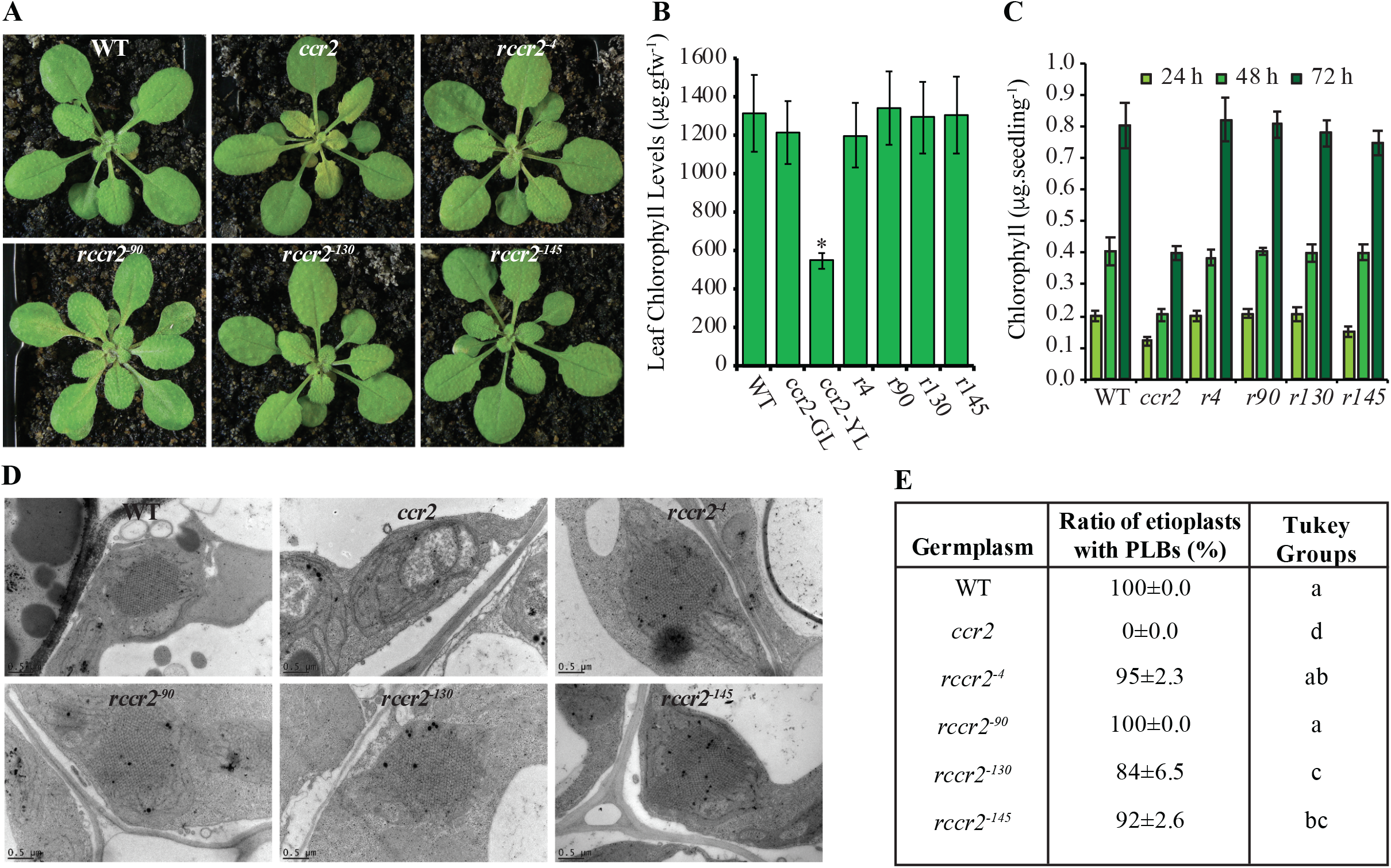
PSY variants restore greening and chlorophyll levels in *ccr2* leaves and cotyledons as well as PLB formation in etiolated seedlings. **A)** Three-week-old WT, *ccr2*, and r*ccr2* mutant variant (r*ccr2*^-4^, r*ccr2*^-90^, r*ccr2*^-130^, r*ccr2*^-145^) plants were grown in multiple independent experiments under a 10-h photoperiod and a representative image from each genotype is displayed. **B**) Total chlorophyll in leaf tissues from WT, *ccr2* and *rccr2* plants growing under 8-h photoperiod. Error bars denote the standard error of the mean (n=5). **C**) Total chlorophyll levels in cotyledons from de-etiolated seedlings (grown in darkness for 4 d) exposed to continuous white light for 3 days and quantified at 0, 24, 48 and 72 h post illumination Error bars denote standard error of the mean (n=20 seedlings). *R*; *rccr2*. Stars denote significance by comparing *rccr2* lines to WT using a one-way ANOVA (*p* < 0.05). **D**) Transmission electron microscopy (TEM) images of representative etioplasts from 5-d old etiolated WT, *ccr2*, and r*ccr2* mutant variant (r*ccr2*^-4^, r*ccr2*^-90^, r*ccr2*^-130^, r*ccr2*^-145^) seedlings. Images are representatives of > 15 plastids from > 5 TEM sections. **E**) Ratios of etioplasts containing PLBs. Letters denote post-hoc Tukey groups following a one-way ANOVA test.

Next-generation sequencing of *rccr2*^−4^, *rccr2*^−90^, *rccr2*^−130^ and *rccr2*^−145^ revealed each harboured a single mutation in the *PSY* (At5g17230) gene (Fig. 2A and Table S2). The single nucleotide polymorphisms (SNPs) were confirmed by Sanger sequencing (Fig. 2B). *rccr2*^−4^ and *rccr2*^−90^ had G➔A mutations at exons 4 and 5 leading to M266I and A352T amino acid changes in PSY respectively referred to as *psy-*4 and psy*-90*. In *rccr2*^−130^ a C➔T mutation at exon 3 resulted in a substitution of P178 to S, referred to as *psy-130*. In *rccr2*^−145,^ the G➔A mutation at the exon 2/intron 3 border generated an alternative splice site that retained intron 3 and potentially introduced a premature stop codon and truncate protein referred to as *psy-145* (Table S2). Albino seedling phenotypes of *rccr2*^−145^ growing in soil were observed (20% of a homozygous population), revealing an atypical segregation of *psy*^−145^ splice variant dominance (Fig. S3A). Both the full-length and truncated versions of the *PSY* transcripts were evident in *rccr2*^−145^, although the full-length transcript was 3-fold higher than the spliced variant in green rosette leaves (Fig. S3B-C). Therefore, the four *psy* gene mutations retain functionality of PSY and only *rccr2*^−145^ displays partial lethality.

**Fig. 2:**
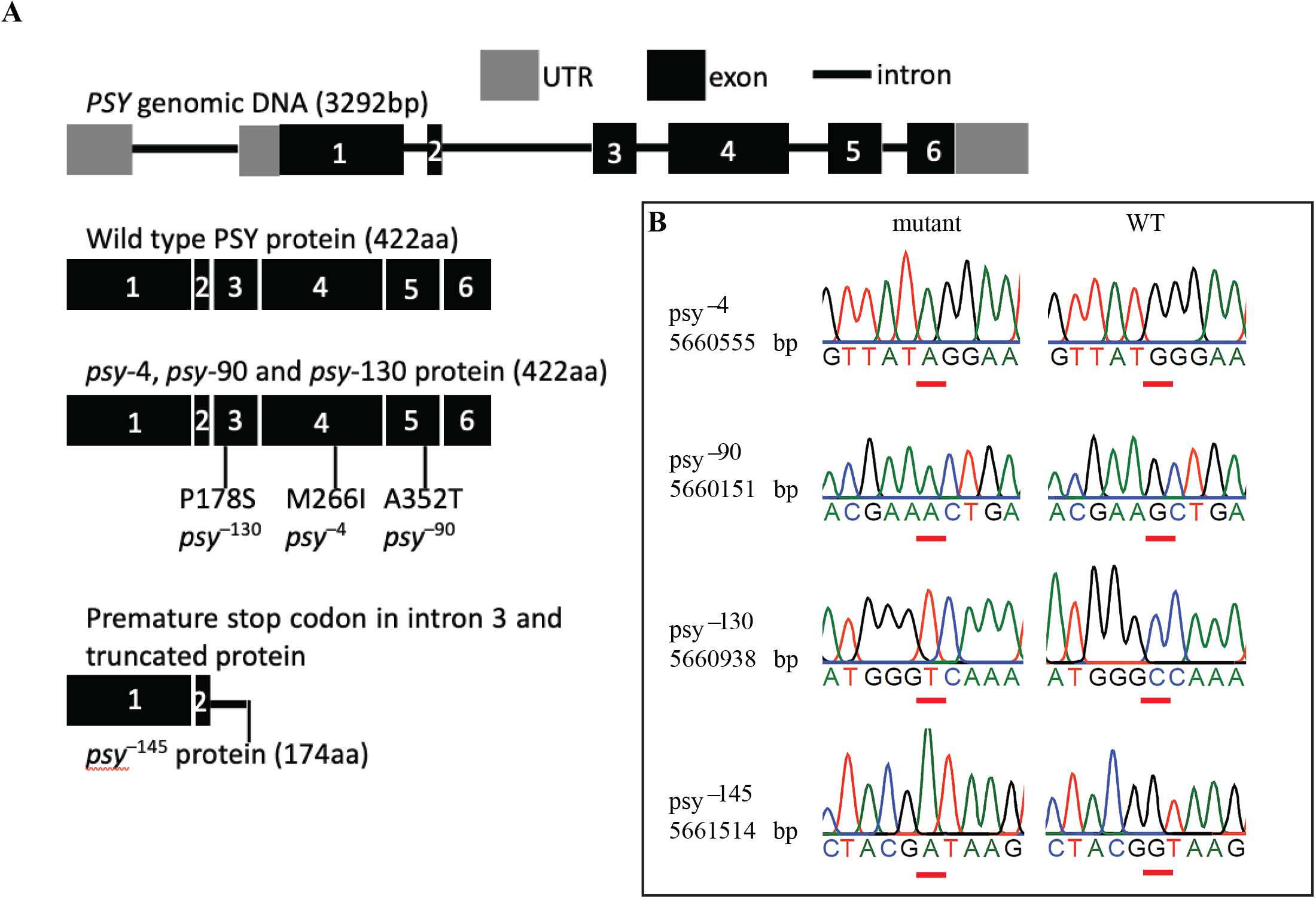
Sequencing of *rccr2* lines identifies mutations within the *PSY* gene that retains protein function. **A**) Schematic structure of wild-type *PSY* DNA, exon structure of the protein and position of the amino acid variant in PSY for r*ccr2*^-4^, r*ccr2*^-90^, and r*ccr2*^-130^. UTR; untranslated leader region, aa; amino acid, P178S; proline at position 178 to serine (r*ccr2*^-130^), M266I; methionine 266 to isoleucine (r*ccr2*^-4^), A352T; alanine 352 to threonine (r*ccr2*^-90^). **B**) Sequencing of *PSY* genomic DNA from leaves of multiple independent plants highlights the mutation position in the four *ccr2 psy* lines.

However, PSY protein levels were reduced in *ccr2 psy* variants relative to WT and *ccr2* (Figure 3A). To verify the PSY casual mutations were responsible for restoring plastid development in *ccr2*, a functional copy of PSY regulated by the CaMV35s promoter was overexpressed (PSY-OE) in the four *psy ccr2* double mutants, *ccr2*, and WT. Western blot analysis confirmed that PSY protein levels in 4-week old foliar tissues were overexpressed at similar levels in WT, *ccr2* and *ccr2 psy* double mutant lines harbouring PSY-OE (Fig. 3A). The overexpression of PSY restored the yellow leaf virescence in all *ccr2 psy*::*PSY*-OE lines grown under 10-h photoperiod (Fig. S4A). Consistent with the yellow leaf (YL) phenotype, a two-fold reduction of total chlorophyll was observed in the YL of *PSY*-OE lines compared to older green leaves from WT or WT::*PSY*-OE lines (Fig. S4B). Chloroplast development appears to have been complemented by genetic perturbations that lower PSY levels, raising questions as to what regulates virescence in *ccr2*.

**Fig. 3:**
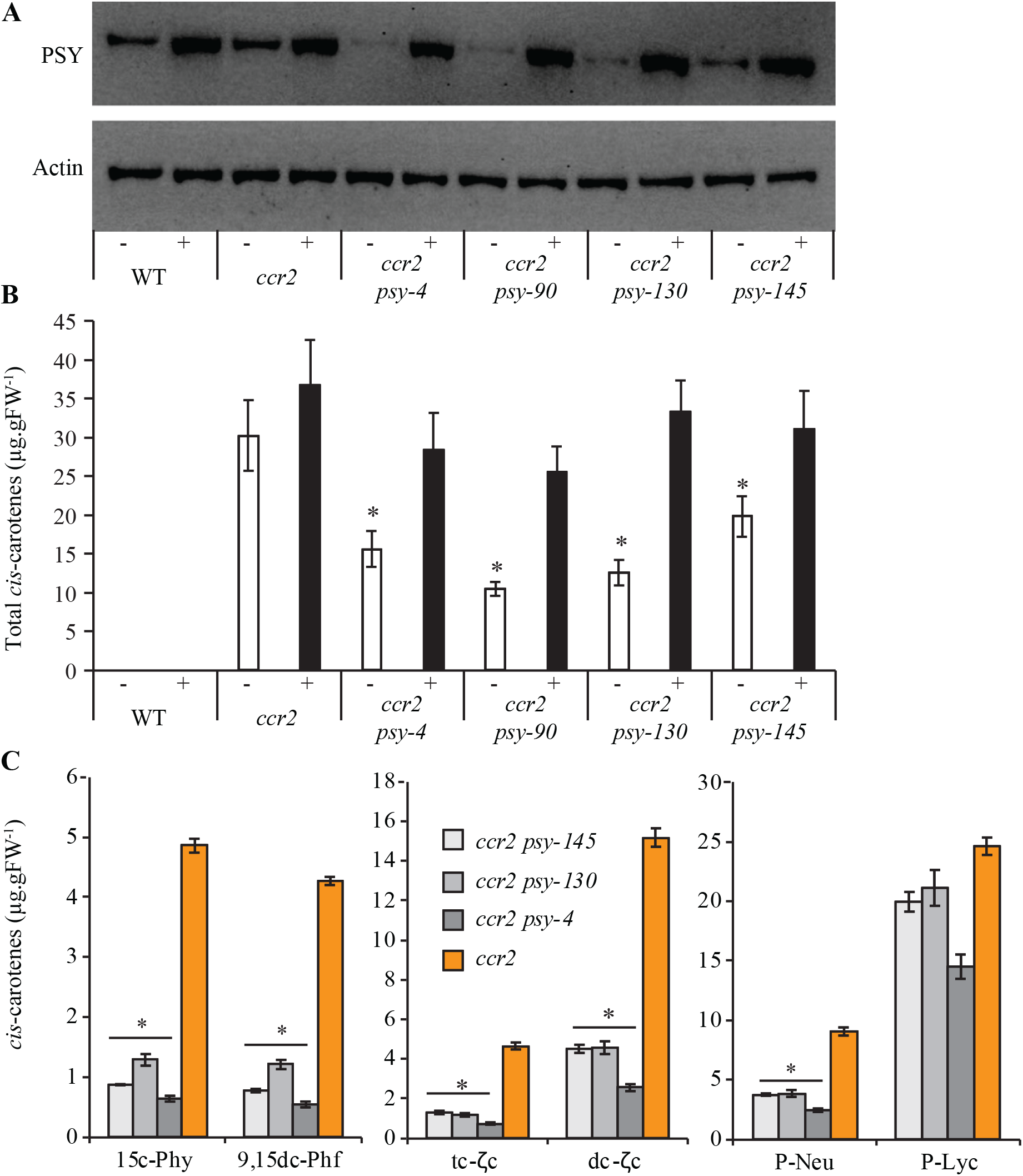
*cis*-carotene and PSY protein levels in WT, *ccr2*, *ccr2 psy*, and *PSY* overexpression lines. **A**) Protein western blot confirming the overexpression of PSY protein in the four *ccr2 psy* double mutants (-) and harboring a transgene overexpressing *PSY* (+). For each sample, 10 µg of total protein extracted from leaf tissues of 4-week-old Arabidopsis plants and the western blots were probed using anti-PSY polyclonal antibody, stripped and re-probed with the ACTIN housekeeper protein. **B**) Total linear *cis*-carotene levels in etiolated (7-d) WT, *ccr2*, and *ccr2 psy* double mutants (-) and harboring a transgene overexpressing *PSY* (+). Error bars denote the standard error of means (n=3), and stars denote significant differences in comparison to *ccr2* (*P* < 0.05; one-way ANOVA). **C**) Linear *cis*-carotene levels in 7-d old etiolated tissues from WT, *ccr2* and *ccr2 psy* double mutants. Error bars denote the standard error of means (n=3). tc-ζ-c: tri-*cis*-ζ-carotene; dc-ζ-c: di-*cis*-ζ-carotene; P-Neu: tri-*cis*-neurosporene; P-Lyc: tetra-*cis*-lycopene; 15c-Phy: 15 *cis*-phytoene; 9,15dc-Phf: 9, 15 di-*cis*-phytofluene.

We quantified linear *cis*-carotene content in etiolated tissues from *ccr2 psy* double mutants (Fig. 3). Total *cis*-carotene content in the *ccr2 psy* was reduced by 40% – 70% relative to *ccr2*. Overexpression of PSY in *ccr2 psy* enhanced total *cis*-carotene content back to *ccr2* levels. *cis*-carotene content in *ccr2* was not affected by *PSY* overexpression, and *cis-*carotenes were not detected in WT (Fig. 3B). Intriguingly, the total *cis*-carotene pool was not significantly different between *ccr2 psy-145* and *ccr2 psy-145* harboring PSY-OE indicating the phenotypes are not proportional to the total *cis*-carotene pool.

We investigated if there is a threshold to the complementation such as an individual *cis*-carotene being present or absent. In comparison to *ccr2*, *ccr2 psy* double mutants (*ccr2 psy-4*, *ccr2 psy-130,* and *ccr2 psy-145*) showed a significant reduction in the levels of phytoene (4 – 6 fold), di-*cis*-phytofluene (4 – 6 fold), tri-*cis*-ζ-carotene (3 – 5 fold), di-*cis*-ζ-carotene (3 – 5 fold) and tri-*cis*-neurosporene (2 fold) (Fig. 3C). The levels of tetra-*cis*-lycopene (prolycopene; P-Lyc) in *ccr2 psy*^−130^ and *ccr2 psy*^−145^ were comparable to *ccr2*, and marginally lower in *ccr2 psy*^−4^ (Fig. 3C). Therefore, the restored plastid biogenesis in *ccr2* by mutations that lower PSY protein levels correlate with a threshold reduction in *cis*-carotenes other than prolycopene.

### psy variant alleles restore PIF3 and HY5 transcript and protein levels in ccr2

We questioned if the *cis*-carotenes had a structural or signalling function by investigating how the *psy* variants regulated *PIF3* and *HY5* in etiolated *ccr2* seedlings. Transcript levels of *PIF3* and *HY5* in *ccr2* were 6-fold higher, and 2-fold lower in *ccr2*, respectively (Cazzonelli *et al*., 2020). In the four *ccr2 psy* mutants, the transcript levels of both *PIF3* and *HY5* were like WT (Fig. 4A). In comparison to WT, PIF3 protein levels were dramatically up-regulated in *ccr2*, yet in the four double mutants, levels were comparable to WT. HY5 protein levels were significantly lower in *ccr2* at trace levels, while all four *ccr2 psy* double mutants displayed considerably higher HY5 protein levels like WT (Fig. 4B). The higher PIF3 to HY5 ratio of transcripts in etiolated *ccr2* seedlings, corroborates with protein levels and the four PSY mutant variants restore WT levels of PIF3 and HY5 levels. Therefore, a threshold level of specific *cis*-carotenes can be linked to the generation of a signalling metabolite.

**Fig. 4:**
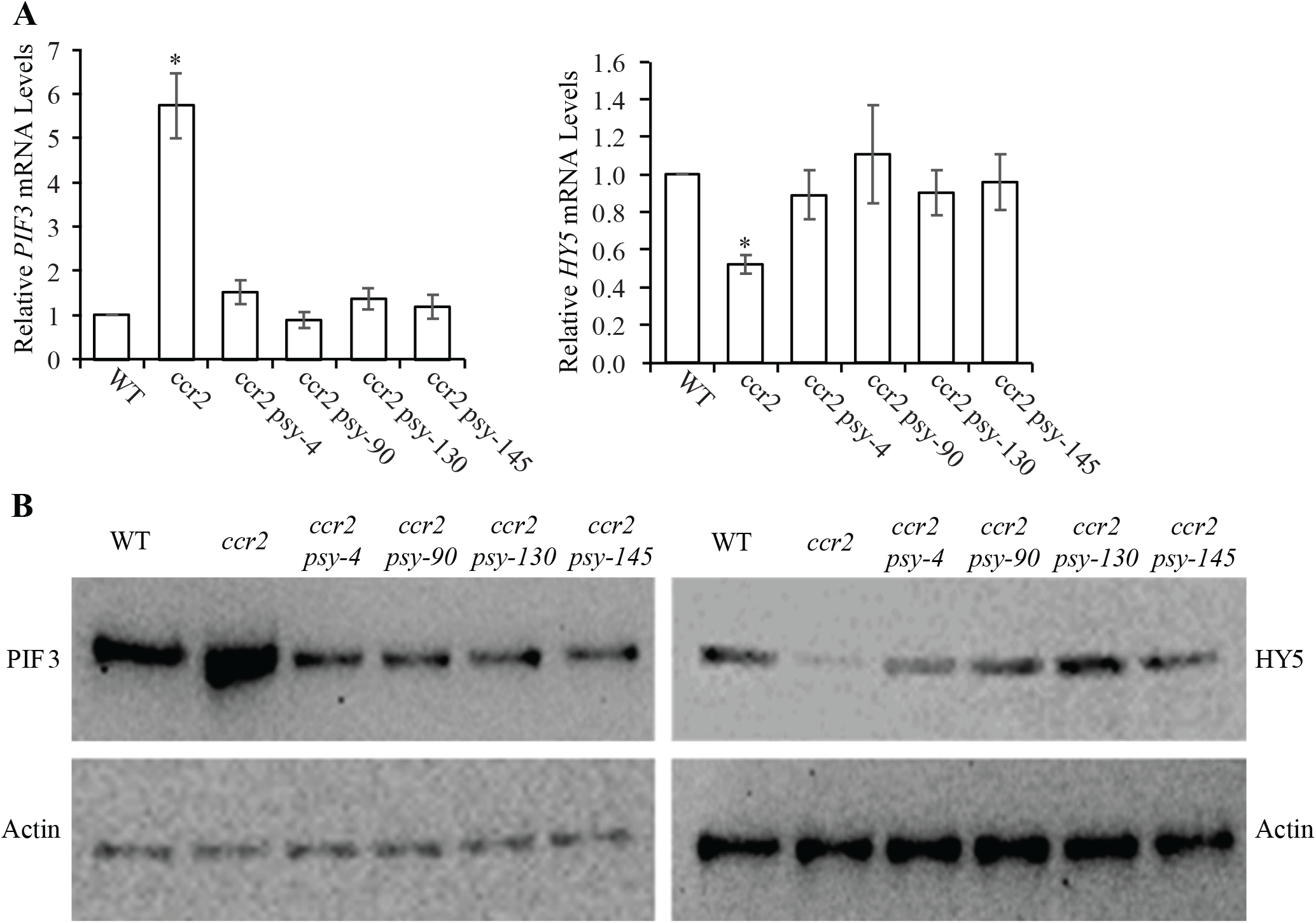
Transcript and protein levels of HY5 and PIF3 in WT, *ccr2* and *ccr2 psy*. **A**) Transcript levels of *PIF3* and *HY5* in etiolated 7-d seedling tissues from WT, *ccr2*, *ccr2 psy-4*, *ccr2 psy-90*, *ccr2 psy-130* and *ccr2 psy-145*. Error bars denote the standard error of the mean (n=3). Stars denote significance by comparing mutant lines to WT using a one-way ANOVA (*p* < 0.05). **B**) Representative western blot images showing protein levels of PIF3 and HY5 in WT, *ccr2* and the *ccr2 psy* etiolated seedling tissues. For each sample, five micrograms of total protein was loaded to separate gels, blotted with PIF or HY5 antibodies and membrane re-probed using anti-Actin antibody as an internal loading control.

### psy variant alleles show reduced PSY enzyme activities

We examined phytoene synthase activity of the four *psy* variant alleles. First, we used an *in vivo* shoot-derived calli (SDC) assay where seeds are germinated in the light to develop seed-derived callus and thereafter transferred to darkness for 14 days (Schaub *et al*., 2018). During darkness ongoing carotenoid biosynthesis is compensated by degradation and norflurazon which inhibits PDS activity enriches for phytoene and reflects the rate of phytoene synthase activity as a proxy for PSY protein levels. SDC of WT and *ccr2* accumulated similar levels of phytoene, which were 1.5-to 3-fold higher than in the four *ccr2 psy* double mutants (Fig. 5A). Endogenous phytoene synthase was purified from WT, *ccr2* and *ccr2 psy* foliar tissues by immunoprecipitation and *in vitro* activity assays showed that PSY variants synthesised less phytoene indicating impaired PSY activity (Fig. 5B). Using an *E. coli* carotenoid expression system generating GGPP (Cunningham and Gantt, 2007), we tested the *in vitro* activity of purified recombinant versions of PSY and its mutant variants (*psy-4*, *psy-90*, *psy-130*) to convert GGPP to phytoene. In agreeance with pervious assays, *psy-4* and *psy-90* displayed significantly reduced phytoene levels, while *psy-130* was marginally lower. Collectively, the *psy* mutant variants showed impaired PSY activity.

**Fig. 5:**
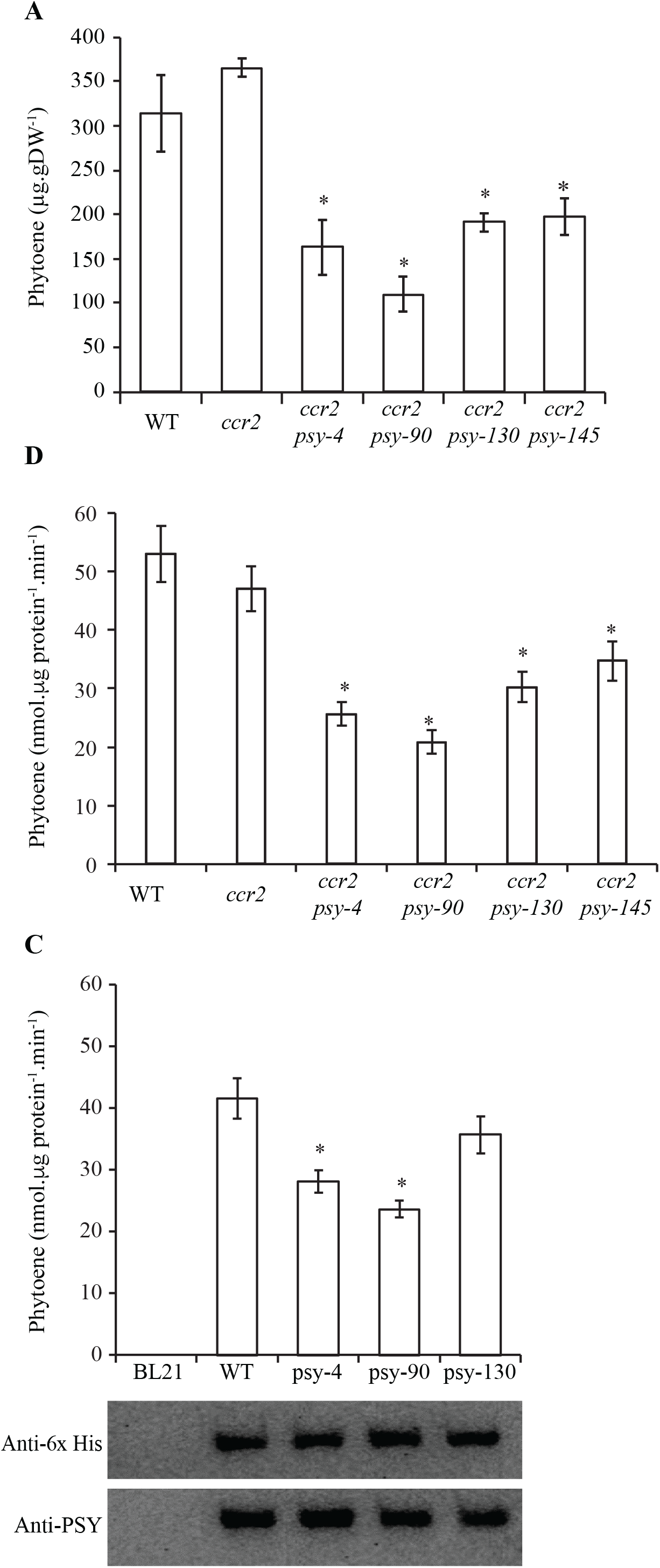
In vivo/vitro enzymatic assays of PSY activity and carotenoid levels in WT, *ccr2* and *ccr2 psy*. **A**) *In vivo* PSY activity assay using seed-derived callus (SDC) treated with 1 µmol.L^-1^ norflurazon. Norflurazon inhibits PDS and leads to the accumulation of phytoene, providing a measure of PSY activity. **B**) *In vitro* activity of endogenous PSY (or mutant) from WT, *ccr2* and *ccr2 psy* double mutants. PSY was purified from mature leaves by co-immunoprecipitation, and 5 μg was used for enzymatic activity assays measuring phytoene levels. **C**) *In vitro* enzyme activity assay showing phytoene and recombinant PSY protein levels purified from *E. coli* cells expressing WT, *psy-4*, *psy-90*, *psy-130* and *psy-145* variants. Untransformed *E. coli* BL21 (DE3) cells were used as a negative control. Western blots showing anti-PSY and anti-6× Histidine (His) levels to confirm expression and purification of recombinant PSY. 5 µg of protein was loaded to the gel. Data is shown as mean ± standard errors of three biological replicates in all measurements. *Difference is significant compared to WT (*P* < 0.05 in one-way ANOVA).

We used computational modelling to predict how the amino acid substitutions of psy-4, psy-90 and psy-130 (Fig. 2A) alter enzyme activity compared to a structural model of PSY (Fig. 6). In PSY-130 a proline to serine substitution (P178S) could impact metal binding at aspartate residues 172 and 176, providing higher flexibility to form a hairpin. The P178S mutation may impact the metal binding by aspartate 302, which is proximal to proline 178 of the predicted 3D structure. These aspartate residues reside within the D_172_ELVD_176_ and D_298_VGED_302_ regions conserved isoprenoid synthase proteins shown to form an active site and bind substrates (Pandit *et al*., 2000). In *psy-4*, the methionine substitution to isoleucine (P266I) is situated close to the lower substrate pocket. Alanine 352, which was mutated to threonine in psy-90 (A353T) models steric interactions with the adjacent α-helix. This could impact substrate binding because of the proximity to the 3D structured substrate pocket (Fig. 6). Therefore, the psy-4 and psy-90 variants likely alter the structure of the active site and/or substrate binding, corroborating with reduced phytoene synthase catalytic activity demonstrated using *in vivo* and *in vitro* assays.

**Fig. 6:**
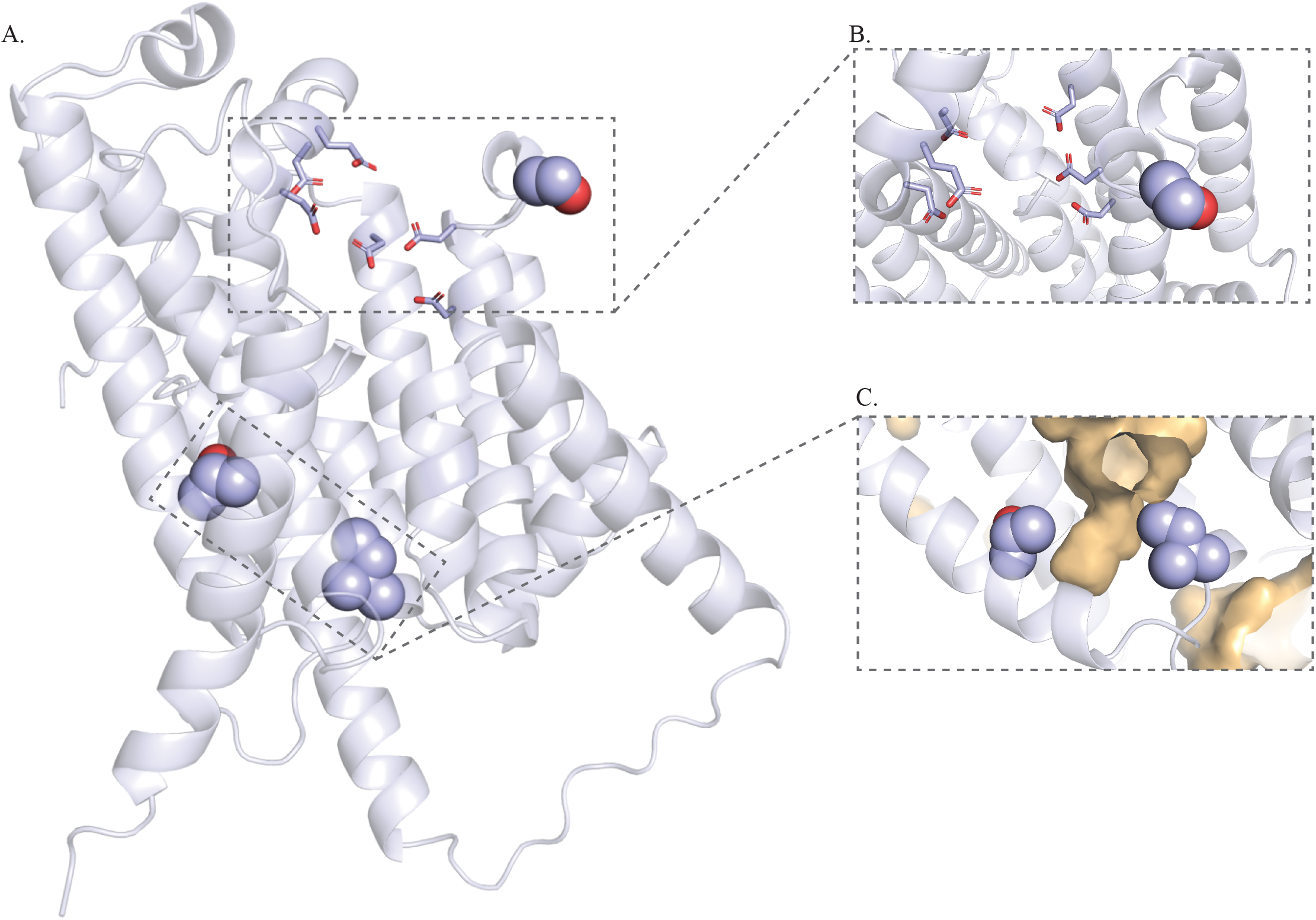
Structural mapping of amino acid substitutions in a 3D PSY structural model. Cartoon representation of a 3D structural model representation built using Colabfold implementation of Alphafold2 and ESMFold. Sub-panels show mutations with different angles and features for clarity. PyMOL cavity settings are Cavities and Pockets (Culled), Cavity Detection Radius at seven angstroms, and Cavity Detection Cutoff at three solvent radii for clarity. **A**) Structural model showing the mutations in ball-representation and putative metal-binding aspartate residues in stick representation. **B**) P178S mutation (shown in ball-representation) in psy-130 and putative metal-binding aspartate residues (shown in stick-representation). This mutation may impact the metal binding by D172 and D176 located in the same hairpin, and that by D302 which is close to P178 in the predicted 3D structure. **C**) A352T (upper left) in psy-90 and M266I (lower right) in psy-4 are shown in ball-representation. Substrate pocket (centre) and voids are shown in tan colour. The A352T substitution may affect substrate binding and the residue’s steric interaction with its adjacent α-helix. The M266I mutation is close to the bottom segment of the substrate pocket.

### psy mutant alleles affect protein-protein interactions between PSY and OR

A split ubiquitin system (SUS) based on yeast-two-hybrid (Y2H) was used to evaluate whether the PSY amino acid substitutions alters protein-protein interaction between PSY and Arabidopsis ORANGE (AtOR) that can post-transcriptionally regulate PSY levels (Zhou *et al*., 2015). A clear interaction was observed between wild-type PSY and AtOR by growth in selective media, and β-galactosidase activity of yeast strains co-expressing different PSY versions fused to C-terminal ubiquitin moiety (Cub) and AtOR fused to N-terminal ubiquitin moiety (Nub) (Fig. 7A). A suppressed interaction was detected when P266I (PSY-4) or A353T (PSY-90), but not P178S (PSY-130), was fused to Cub, while psy-90 generally failed to interact with AtOR (Fig. 7A and 7B).

**Fig. 7:**
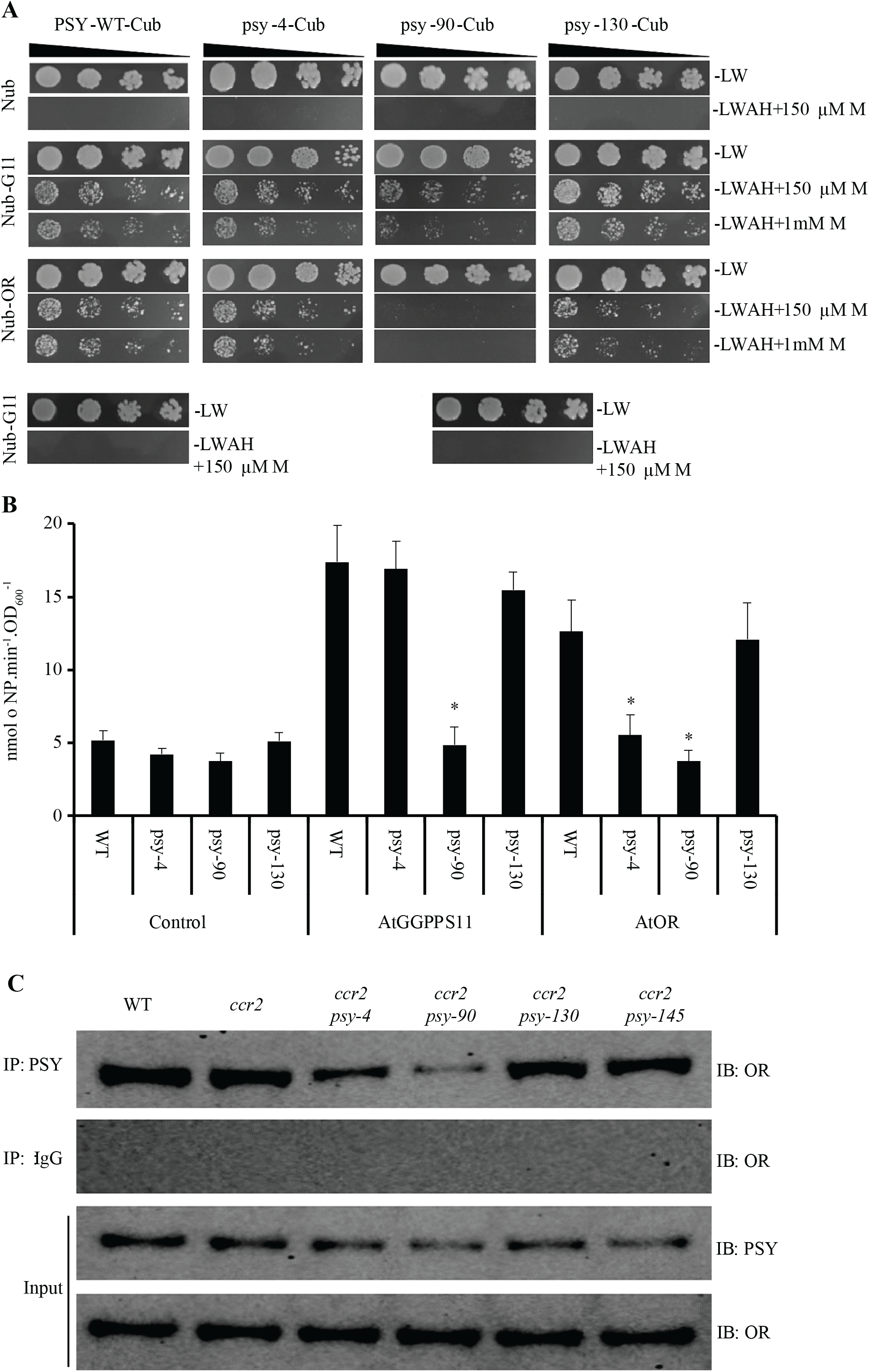
PSY amino acid substitutions alter AtPSY protein-protein interaction with AtOR and AtGGPPS11. **A**) Protein-protein interaction assays using split ubiquitin system. *AtPSY* gene variants were fused to the C-terminal ubiquitin moiety (Cub) and co-expressed with AtOR (OR) or AtGGPPS11 (G11) fused to the N-terminal ubiquitin moiety (Nub). Yeasts were spotted onto either nonselective (-LW) or fully selective medium (-LWAH) in 10-fold dilution series. In order to reduce background activation of reporter genes and visualize different interaction strengths, methionine (150 µmol.L^-1^ and 1 mmol.L^-1^) was added to the media reducing expression of Cub fusion proteins (+150 µM M and +1mM M, respectively). Control combinations with empty Cub expressing vectors are included below. **B**) β-Galactosidase activity of yeast strains co-expressing different PSY versions fused to Cub and AtOR or ArGGPPS11 fused with Nub or Nub only (Control), respectively. Enzyme activity was determined by oNPG assay and is given in nmol oNP.min^-1^.OD_600_^-1^. Error bars indicate standard error of means (n=3). Stars denote significant differences (*p* < 0.05) in comparison to WT (ANOVA). **C**) Co-immunoprecipitation analysis of AtPSY and AtOR proteins. Equal amounts of total protein extracted from each leaf sample were immunoprecipitated using an anti-PSY antibody or normal rabbit IgG as a negative control. Ten microliters of immunoprecipitate was subjected to western blot using anti-OR antibody. Half of the protein before immunoprecipitation was kept for input and subjected to western blotting with anti-PSY and anti-OR antibodies. Experiments were performed in triplicate and representative results were displayed.

We tested the interaction between PSY versions and Arabidopsis GGPP SYNTHASE 11 (AtGGPPS11), which was suggested to be an essential protein interacting with PSY to produce phytoene (Ruiz-Sola *et al*,. 2016). The protein-protein interaction between A353T (PSY-90) and AtGGPPS11 was negatively affected, although P266I (PSY-4) and P178S (PSY-130) variants showed WT interactions with AtGGPPS11 (Fig. 7A and 7B). Consistent with PSY activity assays, the production of phytoene in yeast cells expressing A353T (PSY-90) plus AtGGPPS11 was blocked, while P266I (PSY-4) or P178S (PSY-130) plus AtGGPPS11 showed a -2 to 4-fold reduction in phytoene levels compared to PSY (Fig. S5).

To confirm the reduced interaction between AtOR and PSY variants, we quantified AtOR in the PSY co-immunoprecipitate mixture using western blot analysis. Considerably lower levels of AtOR were observed in the A353T (PSY-90) co-immunoprecipitate, and a clear reduction was evident in the P266I (PSY-4) co-immunoprecipitate evidencing that these two variants, but not P178S (PSY-130) or PSY-140 can impair the physical interaction with AtOR (Fig. 7C).

We next quantified PSY and AtOR protein levels in total protein extracts of WT, *ccr2* and *ccr2 psy* variant leaf tissues. PSY protein levels in *ccr2 psy-4* and *ccr2 psy-90* were lower (Fig. 8), and enzyme-linked immunosorbent assays (ELISAs) confirmed 2- and 5-fold reductions, respectively (Fig. S6). The levels of AtOR protein were also lower in *ccr2 psy-4* and *ccr2 psy-90* by 3-to 4-fold (Fig. 8 and Fig. S6). Western blot and ELISA assays showed that both AtPSY and AtOR protein levels in *ccr2 psy-130* and *ccr2 psy-*145 were like WT (Fig. S6). We concluded that the PSY-AtOR interaction was impaired in *ccr2 psy-4* and *ccr2 psy-90,* leading to a reduction in PSY and AtOR protein levels.

**Fig. 8:**
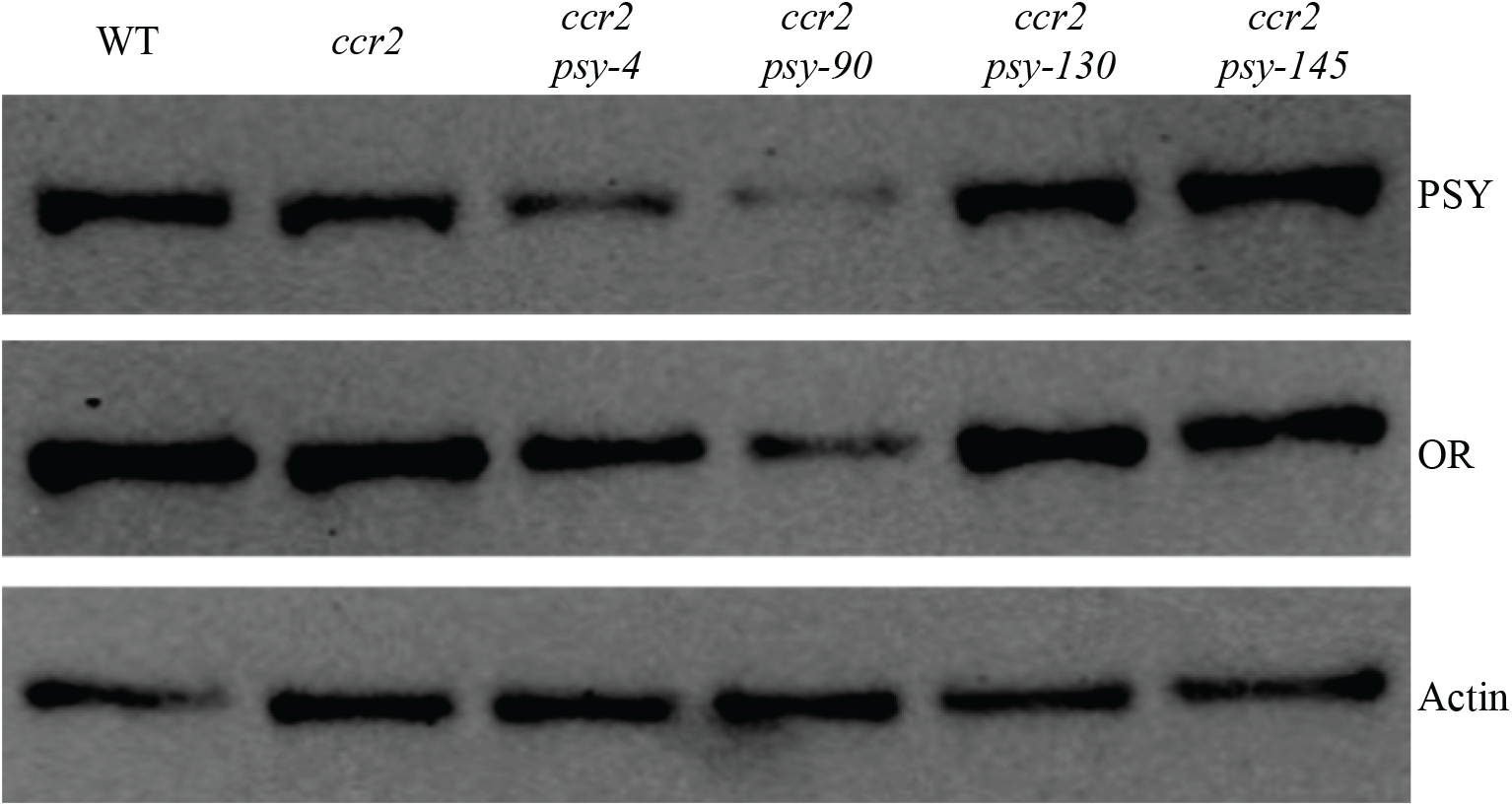
Western blot analysis of PSY and OR protein levels in *ccr2 psy* variants. Ten micrograms of total protein extracted from leaf tissue of WT, *ccr2* or *ccr2 psy* variants was used for western blot analysis to quantify endogenous AtPSY and AtOR protein levels using anti-PSY and anti-OR antibodies, respectively. Actin was included as an internal reference and detected using anti-Actin antibody. The blot shown is representative of three western blot replicates.

### PSY localization was not affected in psy mutant variant alleles

To examine the sub-organellular localization of the PSY versions, we transiently expressed recombinant protein of the PSY versions in protoplasts isolated from cotyledons of Arabidopsis etiolated seedlings and detected PSY or psy variants in PLB as well as stroma fractions of etioplasts. As a control, we included psy*-*N_181_P_270_ (psy-NP), the Arabidopsis equivalent variant of ZmPSY-N_168_P_257_ that was reported to display an altered localization in etioplasts and chloroplasts. PSY takes two topological forms: membrane-bound and stromal, and we detected endogenous PSY in both PLB and stroma fractions (Fig. 9). There was no significant difference among the pattern of distribution of the PSY and PSY variants in the PLB and stroma fractions of etioplasts (Fig. 9). The transiently expressed recombinant PSY or PSY variants appeared to affect PSY levels in the stroma only (Fig. 9).

**Fig. 9:**
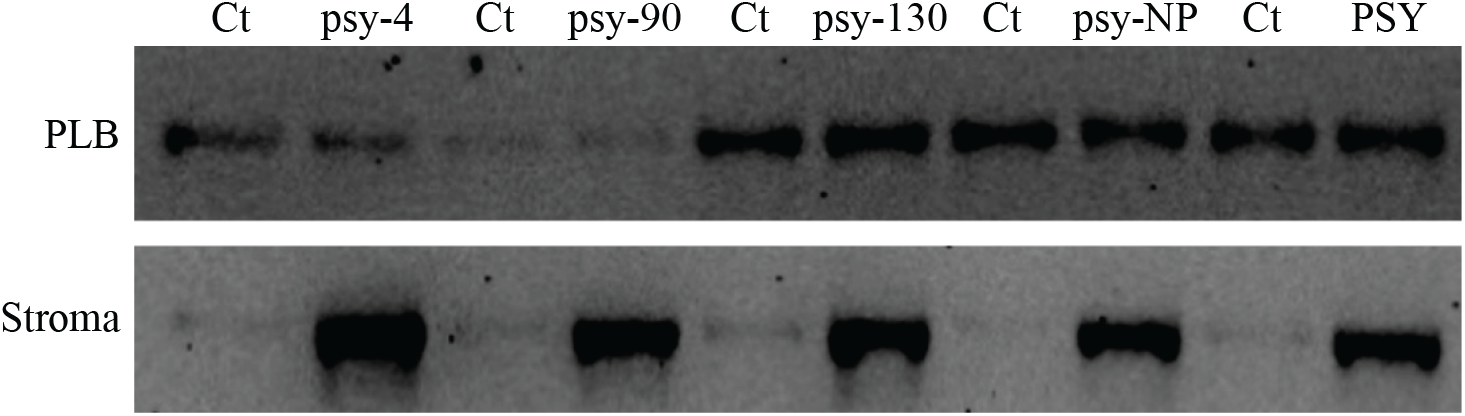
PSY levels in PLB and stroma fractions of Arabidopsis etioplasts before and after the expression of recombinant proteins. Protoplasts were isolated from the cotyledons of etiolated 7-d old seedlings and incubated in the dark with a PSY-CGFP plasmid for 16 h. Ten microliters of PLB or stroma fraction of etioplasts was subjected to western blot using anti-PSY antibody. Ct: control with no expression of recombinant PSY.

## DISCUSSION

Here we define new mutations that disrupt PSY enzyme activity at the entry point to carotenoid biosynthesis that act epistatically to *crtiso* mutations with respect to carotenoid profiles and visual phenotypes, namely impaired leaf greening. A forward genetics screen identified four non-lethal *PSY* mutant alleles harbouring a single point mutation that reduced PSY activity, protein levels, and AtOR interactions, which underpin substrate supply into the carotenoid biosynthetic pathway (Fig. 10). The psy variants significantly reduced threshold levels of specific *cis*-carotenes in *ccr2* and presumably the production of a *cis*-ACS regulating plastid biogenesis and the PIF3/HY5 module (Fig. 10). Single amino acid sequence changes can profoundly alter PSY activity and identification, and optimization of which key amino acid residues reduce activity will help to not only unravel the intrinsic features of PSY activity but facilitate modelling of highly efficient PSY for development of carotenoid enriched crops.

**Fig. 10.**
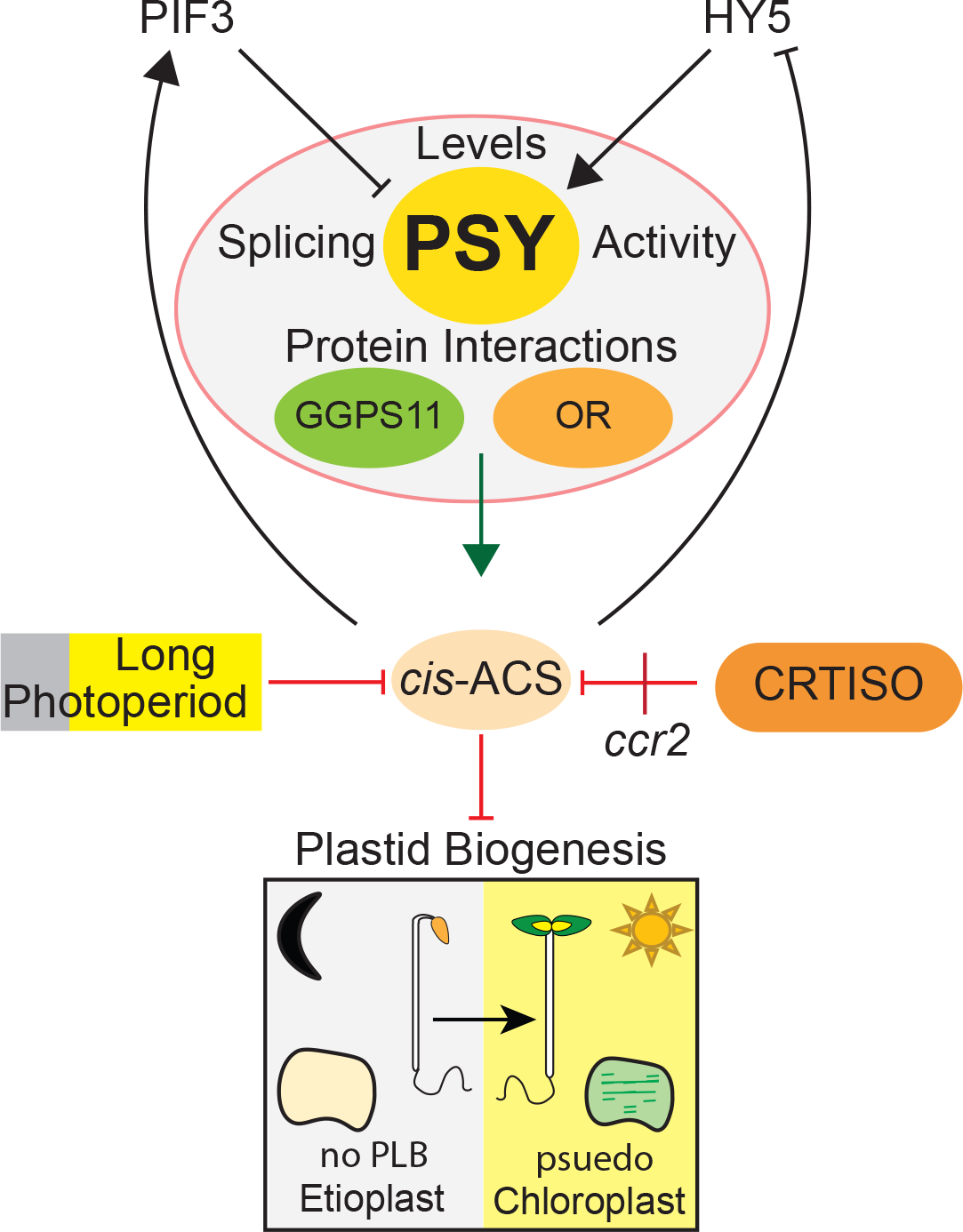
Manipulation of PSY protein activity provides a central target to modify a linear *cis*-carotene-derived apocarotenoid signal (*cis*-ACS) that modulates plastid biogenesis. Variants in PSY activity affect splicing (alternative leading to truncated protein), enzyme activity (enzyme-substrate binding), and protein-protein interactions (multi-enzyme complex binding between PSY, OR, and/or GGPS11) impacting upon protein levels that constitute dynamic regulations to fine-tune the level of a *cis*-ACS without impacting overall carotenoid end-product accumulation. *cis*-ACS up-regulates PIF3 and suppresses HY5 protein levels that fine-tune PSY control over *cis*-ACS biosynthesis. CRTISO activity and longer photoperiods (i.e., efficient photoisomerization) reduce *cis*-carotene biosynthesis and the Arabidopsis *crtiso* loss-in-function *ccr2* mutant causes *cis*-carotene accumulation that triggers *cis*-ACS to block prolamellar body (PLB) formation in etioplasts and causes psudeochloroplasts to form when mutants are grown under shorter photoperiods (Cazzonelli *et al*., 2020).

### Mutations in PSY alter RNA splicing, enzyme activity, protein levels, and multi-enzyme interactions

Mechanisms comprised of alternative splicing, enzyme-substrate interactions, multi-enzyme complex binding, constitute dynamic regulation to fine-tune PSY activity and reduce linear *cis*-carotene biosynthesis without impacting overall carotenoid end-product accumulation. Single amino acid changes in PSY have been shown to increase enzymatic activity and up-regulate carotenoid biosynthesis (Shumskaya *et al*., 2012; Welsch *et al*., 2010). Here we defined four unique mutations that impaired PSY operations in *ccr2,* leading to the restoration of plastid biogenesis (Fig. 1). Alternative splicing changes in *PSY* transcript sequence length have been shown to produce variants with different translation efficiency and/or distinct enzyme activity to control the functional PSY in Arabidopsis leaves (Alvarez *et al*., 2016; Zhou *et al*., 2022). The mutation in *psy-* 145 altered intron-3 splicing leading to a truncated protein evident by 20% of *ccr2 psy-145* homozygous seedlings displaying an albino phenotype (Fig. 2, S2, Table S2). The remaining 80% of *ccr2 psy-145* seedlings remained green, harbouring both the spliced and non-spliced variants of PSY, retained interactions between PSY and AtOR without affecting their protein levels (Fig. 3, 6, 7 and 8), yet reduced phytoene levels *in vivo* and *in vitro* activity assays (Fig 5). The reduced phytoene levels in *ccr2 psy-145* compared to *ccr2* could be due to the inefficiency in intron-3 splicing (Fig. S3), keeping consistent with the reduction in *cis*-carotene levels in *ccr2 psy-145* (Fig. 3).

Three non-lethal mutations impaired PSY operations by reducing PSY activity for *psy-*4, *psy-* 90 and *ps y*1*-*30 when tested *in vivo* (SDC) and *in vitro* (*E.coli* assays) (Fig 5). The amino acid substitutions M266I (psy-4), A352T (psy-90), P178S (psy-130) modified PSY operations to lower accumulation and *cis*-carotene in *ccr2* (Fig. 3). The AtPSY _172_DELVD_176_ and _298_DVGED_302_ sequences were predicted to form an active site and bind phosphate groups of GGPP, and are conserved among isoprenoid synthase superfamily (Pandit *et al*., 2000; Shumskaya *et al*., 2012; Zhou *et al*., 2022). Our modelling of the PSY enzyme showed that these two sequences were positioned at the entry to the substrate pocket (Fig. 6). The P178S mutation of psy-130 (Fig. 2, Table S2) is between the _172_DELVD_176_ sequence and S_181,_ which is an essential amino acid for the activity and localization of PSY (Shumskaya *et al*., 2012). P178S could alter the structure of the substrate pocket and/or impact putative metal binding at D172, D176 and D302 within the two conserved PSY protein domains (Fig. 6). The M266I mutation in psy-4 is close to the bottom segment of the substrate pocket and likely effects substrate fitting and/or reaction catalysis. The A352T mutation of psy-90 is also close to the substrate pocket (Fig. 6) and may interfere with the formation of a preferable structure of the pocket. The transient expression of psy-4, psy-90, psy-130, and Atpsy*-*N_181_P_270_ (psy-NP) variants in Arabidopsis protoplasts showed similar localisation in PLB and stroma fractions when compared to the WT PSY (Fig. 9). While Atpsy*-*N_181_P_270_ can be localised to plastoglobuli in rice and maize protoplasts (Shumskaya *et al*., 2012) this was not the case in Arabidopsis. The P178S (psy-130), M266I (psy-4), and A352T (psy-90) mutations do not affect plastid localisation yet impair PSY enzyme-substrate interactions that reduces *cis*-carotene biosynthesis.

The PSY:AtOR interaction was reduced in *psy-*4 and *psy-*90 (Fig 7, 8, 9), thereby reducing the levels of both proteins and phytoene levels in their respective *ccr2 psy* variant backgrounds (Fig. 5A, 5B, 8, S6). The reduced PSY:AtOR interaction and lower protein levels are consistent with a previous report (Zhou *et al*., 2015), where a decrease in AtOR protein levels negatively affected the recruitment of inactive PSY populations from the stroma to the membrane and consequently lowered PSY activity. Furthermore, reduced interaction between PSY and OR increases the proportion of PSY which is subjected to Clp-mediated degradation, thus reducing the overall amount of enzymatically active PSY (Welsch *et al*., 2018). A nonsense mutation in *Cucumis melo* OR protein (*Cmor*-lowβ) was also able to lower tetra-*cis*-lycopene levels in *crtiso* melon fruit tissues (Chayut *et al*., 2017). Therefore, the impairment in PSY-AtOR interaction in *ccr2 psy-4* and *ccr2 psy-90* leads to a reduction in PSY and AtOR protein levels reveals a new rate-limiting step to modulate the biosynthesis of *cis*-carotenes and an associated signalling metabolite.

For the biosynthesis of carotenoids, a large multi-enzyme complex appears to involve PSY and GGPPS (Camara, 1993; Cunningham and Gantt, 1998; Fraser *et al*,. 2000; Shumskaya and Wurtzel, 2013). Within the metabolon, GGPPS may channel its product GGPP to PSY as substrate; hence, interaction with GGPPS is required for PSY activity (Ruiz-Sola *et al*,. 2016). The A352T mutation in psy-90 perturbed the PSY-AtGGPPS11 interaction such that in yeast cells phytoene synthesis was almost abolished evidencing that full PSY activity requires interaction with GGPPS11 (Fig. 7 and S5). Our study herein demonstrates that PSY, AtOR and GGPPSS11 constitute a multi-enzyme complex that acts to bottleneck the supply of linear *cis-*carotene substrates required to generate a *cis*-ACS that regulates plastid development (Fig. 10).

### psy variants reduce di-cis-ζ-carotene and tri-cis-neurosporene substrates

Acyclic linear *cis*-carotene levels in *ccr2 psy* variants were greatly reduced below the threshold in *ccr2* that signals control over plastid biogenesis. The psy variants reduced total linear *cis*-carotene levels in etiolated seedlings by 2-to 3-fold, yet there was only a significant reduction in phytoene, di-*cis*-phytofluene, tri-*cis*-ζ-carotene, di-*cis*-ζ-carotenes and tri-*cis*-neurosporene, but not tetra-*cis*-lycopene, ruling the later out as a substrate for ACS (Fig 3, 4). We rule out phytoene, phytofluene, and tri-*cis*-ζ-carotene as potential substrates for generating a *cis*-ACS since the *ccr2 ziso* double mutants continue to accumulate these substrates (Cazzonelli *et al*., 2020) and just like *ccr2 psy* variants show restored plastid biogenesis. Evidence using chemical inhibition of CCD activity supported that tri-*cis*-neurosporene and di-*cis*-ζ-carotene were potential substrate(s) for *in vivo* cleavage into a signalling metabolite (Cazzonelli *et al*., 2020). Collectively both these studies reveal that di-*cis*-ζ-carotene and/or tri-*cis*-neurosporene are the preferred substrate(s) of a *cis*-ACS (Fig. 10).

### psy variants modulate the PIF3/HY5 regulatory hub

PIF3 and HY5 are key regulatory transcription factors controlling the dark-to-light transition of plants in concert with carotenogenesis. A high PIF3/HY5 ratio maintains skotomorphogenesis in etiolated seedlings, whereas a low PIF3/HY5 ratio promotes photomorphogenesis in de-etiolated seedlings (Dong *et al*., 2014; Job and Datta, 2020; Osterlund *et al*., 2000). PIF3 and HY5 negatively and positively regulate *PSY* expression, respectively, affecting carotenoid biosynthesis (Fig. 10) (Toledo-Ortiz *et al*., 2010; Toledo-Ortiz *et al*., 2014). DET1 and constitutive photomorphogenic 1 (COP1) modulate the PIF3/HY5 regulatory hub by post-transcriptionally controlling the levels of PIF3 and HY5 (Lau and Deng, 2012; Llorente *et al*., 2017; Stephenson *et al*., 2009; Xu *et al*., 2016). Using a chemical inhibitor of CCD cleavage, we previously showed that a *cis*-ACS can act independent of DET1 to transcriptionally regulate *PIF3* and *H Y 5*mRNA expression and their corresponding protein levels (Cazzonelli *et al*., 2020). Here we evidenced that limiting a threshold level of specific *cis*-carotenes in *ccr2* can restore the ratio of PIF3 and HY5 in dark grown seedlings such that it mimics a WT plant. Interestingly, the modulation of a higher PIF3/HY5 protein ratio (ie., transcript and protein levels of *PIF3* were up-regulated, while *HY5* was down-regulated) does not affect *PSY* protein levels in *ccr2* etiolated tissues. Rather the modification of PSY protein levels/activity in the four *ccr2 psy* mutants can lower the PIF3/HY5 ratio by decreasing *PIF3* and increasing *HY5* transcript and protein levels (Fig. 4). The reduction of PSY protein/activity levels in *psy* variants likely fine-tunes threshold levels of a *cis*-carotene derived signal that regulates the PIF3/HY5 module without affecting end-product carotenoid accumulation.

In summary, a lack of carotenoid isomerisation and shorter photoperiod can trigger threshold levels of *cis*-carotenes that signal transcriptional regulation of the PIF3/HY5 module to control plastid biogenesis during both skotomorphogenesis and photomorphogenesis. Non-lethal point mutations in four *psy* genetic variants manipulated PSY alternative splicing, activity, PSY-AtOR interactions, and PSY protein levels reduced levels of di-*cis*-ζ-carotene and/or tri-*cis*-neurosporene, presumably below a threshold required for ACS production. Given these are the most likely substrates for a yet-to-be-identified *cis*-ACS it remains of interest as to the physiological and biochemical basis of this threshold, which are more than quantities normally required for the production of a hormone. The epistasis between CRTISO and PSY is paramount to controlling *cis*-carotene signalling in plants when light levels are limiting (Fig. 10).

## Acknowledgements

The Authors thank Julian Koschmieder, Dennis Schlossarek and Carmen Schubert from the Faculty of Biology II, University of Freiburg, D79104 Freiburg, Germany for their technical support to quantify pigments in shoot derived calli. We thank Jiwon Lee (Centre for Advanced Microscopy, The Australian National University, Canberra, ACT 2601, Australia) for technical support in Transmission electron microscopy. We thank John Rivers, Kai Xun Chan (The Australian National University), and Dr Ryan McQuinn (Western Sydney University) for insightful discussions.

## Author contributions

Xin Hou; Data curation, Software, Formal analysis, Validation, Investigation, Methodology, Writing (original draft), review, and editing, Prepared figures and tables, Performed most experiments.

Christopher I Cazzonelli; Conceptualization, Resources, Formal analysis, Supervision, Funding acquisition, Validation, Investigation, Visualization, Methodology, Project administration, Performed preliminary experiments, Figure preparation, Writing, review, and editing, Supervised YA and XH.

Ralf Welsch and Julian Koschmieder; Data curation, Formal analysis, Investigation, Methodology, review, and editing, Contributed to Fig. 5.

Yagiz Alagoz; Data curation, Formal analysis, Investigation, Methodology, review, and editing, Contributed Fig. 3 and 10.

Matthew D Mortimer; Investigation, Methodology, Writing – method for protein modelling, review, and editing, Contributed to Fig. 6.

Barry J Pogson; Conceptualization, Resources, Formal analysis, Supervision, Funding acquisition, Validation, Investigation, Visualization, Methodology, Project administration, Writing, review, and editing, Supervised XH.

## Data availability

All data supporting the findings of this study are available within the paper and within its supplementary materials published online. The Arabidopsis Genome Initiative locus number for the major gene discussed in this article is as following: *PSY* (At5g17230). The model of *PSY* gene is drawn from At5g17230.1.

## SUPPLEMENTAL FIGURES

**Supplemental Fig. 1: Simplified diagram of the carotenoid biosynthesis pathway**. Genes encoding the catalysing enzymes of each step before all-*trans*-lycopene are labelled to the left of the pathway and mutants to the right (red line indicates blocked step). GGPP: geranylgeranyl diphosphate; PSY: phytoene synthase; phytoene: *cis*-phytoene; phytofluene: di-*cis*-phytofluene; PDS: phytoene desaturase; ZISO: 15-*cis*-ζ-carotene isomerase; ZDS: ζ-carotene desaturase; CRTISO: carotenoid isomerase. Norflurazon is a herbicide that inhibits PDS activity. *cis*-ACS: linear *cis*-carotene derived apocarotenoid signal (ACS).

**Supplemental Fig. 2: Percentage of leaf virescence leaves in *rccr2* lines reverted the leaf-yellowing phenotype in *ccr2* and led to albino phenotype in one line. A**) Percentage of yellow leaf area of WT, *ccr2* and *rccr2* (r) lines. Plants were grown under 8-h photoperiod. Star denotes a significant difference compared to WT (*p* < 0.05 in one-way ANOVA).

**Supplemental Fig. 3: Characterisation of *ccr2 psy-145* and alternative splicing of *PSY*. A)** Albino seedlings displayed by *rccr2*^−145^ *(ccr2 psy-145* **-** red circle) seedlings were grown under a 16-h photoperiod. **B**) Reverse transcription PCR (RT-PCR) showing alternative splicing of intron 3 in *ccr2 psy*^−145^ leaf tissues. Exons 2 and 3 were partially amplified, and products were visualised on a 1% agarose gel. Spliced shows 221 bp (WT and *ccr2*) and unspliced 762 bp (*ccr2 psy* ^−145^) amplicons. 100 bp ladder is displayed. **C**) Quantitative RT-PCR (qRT-PCR) of *PSY* mRNA amplifying part of intron 3 (162 bp) in *ccr2 psy* ^−145^ (+) or a region (155bp) spanning exon 2 and 3 when intron 3 is spliced out in WT and *ccr2* leaf tissues. Standard error bars are shown (n=10). RNA levels were normalized to Protein Phosphatase 2A (AT1G13320) reference gene validated for mRNA normalisation in leaves using Cyclophilin (At2g29960) and TIP41 (At4g34270) secondary reference genes (Cazzonelli *et al*., 2014; Cazzonelli *et al*., 2010b).

**Supplemental Fig. 4: Overexpression of functional *PSY* restores *ccr2* mutant phenotypes in *ccr2 psy* double mutant lines. A)** Representative images of rosettes show yellow virescence in newly emerged leaves from *ccr2* and *ccr2 psy* lines compared to the WT control and WT::*PSY*-OE. T4 generation transgenic plants were grown under a 10-h photoperiod for 3 weeks, and images from 50-100 plants were analysed for five independent lines. *PSY*-OE; *PSY* overexpression. **B**) Total chlorophyll in leaves from T4 generation transgenic plants. Plants were grown under an 8-h photoperiod. Error bars denote the standard error of means (n=5). Star denotes a significant difference compared to WT-OE (*p* < 0.05 in one-way ANOVA). YL; yellow leaf.

**Supplemental Fig. 5: Split ubiquitin assays showing phytoene levels generated in yeast cells expressing PSY+AtGGPPS11 combinations.** Phytoene levels were measured using HPLC. Error bars indicate standard error of means (n=3). Star denotes a significant difference compared to WT *(p* < 0.05; one-way ANOVA).

**Supplemental Fig. 6: ELISA of PSY and AtOR protein levels.** Each well was coated with total protein from Arabidopsis leaf tissue and incubated with anti-PSY or anti-OR antibodies. Following the addition of substrate solution containing O-phenylenediamine dihydrochloride and H_2_O_2_ absorbances were measured at 492 nm. Values were averaged from five biological replicates, and error bars denote the standard error of means (n=5). Stars denote the significant difference compared to WT (*p* < 0.05; one-way ANOVA).

## SUPPLEMENTAL TABLES

**Supplemental Table 1: Primers used in this study**

**Supplemental Table 2: Genomic information of the mutations identified in *PSY* from *rccr2* lines.** SNP; single nucleotide polymorphism.

## REFERENCES

Alagoz Y, Dhami N, Mitchell C, Cazzonelli CI . 2020. cis/trans Carotenoid Extraction, Purification, Detection, Quantification, and Profiling in Plant Tissues. Methods Mol Biol 2083, 145–163.

Alagoz Y, Nayak P, Dhami N, Cazzonelli CI . 2018. cis-carotene biosynthesis, evolution and regulation in plants: The emergence of novel signaling metabolites. Arch Biochem Biophys 654, 172–184.

Alvarez D, Voss B, Maass D, Wuest F, Schaub P, Beyer P, Welsch R . 2016. Carotenogenesis Is Regulated by 5 ’ UTR-Mediated Translation of Phytoene Synthase Splice Variants. Plant Physiology 172, 2314–2326.

Anwar S, Brenya E, Alagoz Y, Cazzonelli CI . 2021. Epigenetic Control of Carotenogenesis During Plant Development. Critical Reviews in Plant Sciences 40, 23–48.

Anwar S, Nayak J, Alagoz Y, Wojtalewicz D, Cazzonelli CI. 2022. Purification and use of carotenoid standards to quantify cis-trans geometrical carotenoid isomers in plant tissues. In: Wurtzel ET, ed. Methods in Enzymology Carotenoids: Carotenoid and apocarotenoid analysis: Elsevier.

Austin RS, Vidaurre D, Stamatiou G, Breit R, Provart NJ, Bonetta D, Zhang J, Fung P, Gong Y, Wang PW, McCourt P, Guttman DS . 2011. Next-generation mapping of Arabidopsis genes. Plant J 67, 715–725.

Avendano-Vazquez AO, Cordoba E, Llamas E, San Roman C, Nisar N, De la Torre S, Ramos-Vega M, Gutierrez-Nava MD, Cazzonelli CI, Pogson BJ, Leon P . 2014. An Uncharacterized Apocarotenoid-Derived Signal Generated in zeta-Carotene Desaturase Mutants Regulates Leaf Development and the Expression of Chloroplast and Nuclear Genes in Arabidopsis. Plant Cell 26, 2524–2537.

Baranski R, Cazzonelli CI . 2016. Carotenoid biosynthesis and regulation in plants. In: Kaczor A, Baranska M, eds. Carotenoids: Nutrition, Analysis and Technology. Hoboken: Wiley-Blackwell, 161–189.

Camara B . 1993. Plant Phytoene Synthase Complex - Component Enzymes, Immunology, and Biogenesis. Methods in Enzymology 214, 352–365.

Cao H, Zhang J, Xu J, Ye J, Yun Z, Xu Q, Deng X . 2012. Comprehending crystalline beta-carotene accumulation by comparing engineered cell models and the natural carotenoid-rich system of citrus. J Exp Bot 63, 4403–4417.

Cazzonelli CI . 2011. Carotenoids in nature: insights from plants and beyond. Functional Plant Biology 38, 833–847.

Cazzonelli CI, Hou X, Alagoz Y, Rivers J, Dhami N, Lee J, Marri S, Pogson BJ . 2020. A cis-carotene derived apocarotenoid regulates etioplast and chloroplast development. Elife 9, e45310.

Cazzonelli CI, Nisar N, Hussain D, Carmody ME, Pogson BJ . 2010a. Biosynthesis and Regulation of Carotenoids in Plants-Micronutrients, Vitamins and Health Benefits. Plant Developmental Biology: Biotechnological Perspectives Vol 2, 117–137.

Cazzonelli CI, Nisar N, Roberts AC, Murray KD, Borevitz JO, Pogson BJ . 2014. A chromatin modifying enzyme, SDG8, is involved in morphological, gene expression, and epigenetic responses to mechanical stimulation. Front Plant Sci 5, 533.

Cazzonelli CI, Roberts AC, Carmody ME, Pogson BJ. 2010b. Transcriptional control of SET DOMAIN GROUP 8 and CAROTENOID ISOMERASE during Arabidopsis development. Mol Plant 3, 174–191.

Chayut N, Yuan H, Ohali S, Meir A, Sa’ar U, Tzuri G, Zheng Y, Mazourek M, Gepstein S, Zhou X, Portnoy V, Lewinsohn E, Schaffer AA, Katzir N, Fei Z, Welsch R, Li L, Burger J, Tadmor Y . 2017. Distinct Mechanisms of the ORANGE Protein in Controlling Carotenoid Flux. Plant Physiol 173, 376–389.

Cheminant S, Wild M, Bouvier F, Pelletier S, Renou JP, Erhardt M, Hayes S, Terry MJ, Genschik P, Achard P . 2011. DELLAs regulate chlorophyll and carotenoid biosynthesis to prevent photooxidative damage during seedling deetiolation in Arabidopsis. Plant Cell 23, 1849–1860.

Clough SJ, Bent AF . 1998. Floral dip: a simplified method for Agrobacterium-mediated transformation of Arabidopsis thaliana. Plant J 16, 735–743.

Cunningham FX, Gantt E. 1998. Genes and Enzymes of Carotenoid Biosynthesis in Plants. Annu Rev Plant Physiol Plant Mol Biol 49, 557–583.

Cunningham FX, Jr., Gantt E . 2007. A portfolio of plasmids for identification and analysis of carotenoid pathway enzymes: Adonis aestivalis as a case study. Photosynth Res 92, 245–259.

Czechowski T, Stitt M, Altmann T, Udvardi MK, Scheible WR . 2005. Genome-wide identification and testing of superior reference genes for transcript normalization in Arabidopsis. Plant Physiol 139, 5–17.

Dhami N, Pogson BJ, Tissue DT, Cazzonelli CI . 2022. A foliar pigment-based bioassay for interrogating chloroplast signalling revealed that carotenoid isomerisation regulates chlorophyll abundance. Plant Methods 18, 18.

Dong J, Tang D, Gao Z, Yu R, Li K, He H, Terzaghi W, Deng XW, Chen H . 2014. Arabidopsis DE-ETIOLATED1 represses photomorphogenesis by positively regulating phytochrome-interacting factors in the dark. Plant Cell 26, 3630–3645.

Earley KW, Haag JR, Pontes O, Opper K, Juehne T, Song K, Pikaard CS . 2006. Gateway-compatible vectors for plant functional genomics and proteomics. Plant J 45, 616–629.

Escobar-Tovar L, Sierra J, Hernández-Muñoz A, McQuinn RP, Mathioni S, Cordoba E, Colas des Francs-Small C, Meyers BC, Pogson. B, Loen P. 2021. Deconvoluting apocarotenoid-mediated retrograde signaling networks regulating plastid translation and leaf development. Plant J 105, 1582–1599.

Fraser PD, Enfissi EM, Halket JM, Truesdale MR, Yu D, Gerrish C, Bramley PM. 2007. Manipulation of phytoene levels in tomato fruit: effects on isoprenoids, plastids, and intermediary metabolism. Plant Cell 19, 3194–3211.

Fraser PD, Schuch W, Bramley PM . 2000. Phytoene synthase from tomato (Lycopersicon esculentum) chloroplasts - partial purification and biochemical properties. Planta 211, 361–369.

Hartwig B, James GV, Konrad K, Schneeberger K, Turck F . 2012. Fast isogenic mapping-by-sequencing of ethyl methanesulfonate-induced mutant bulks. Plant Physiol 160, 591–600.

Hou X, Rivers J, Leon P, McQuinn RP, Pogson BJ . 2016. Synthesis and Function of Apocarotenoid Signals in Plants. Trends Plant Sci 21, 792–803.

Job N, Datta S . 2020. PIF3/HY5 module regulates BBX11 to suppress protochlorophyllide levels in dark and promote photomorphogenesis in light. New Phytol.

Jumper J, Evans R, Pritzel A, Green T, Figurnov M, Ronneberger O, Tunyasuvunakool K, Bates R, Zidek A, Potapenko A, Bridgland A, Meyer C, Kohl SAA, Ballard AJ, Cowie A, Romera-Paredes B, Nikolov S, Jain R, Adler J, Back T, Petersen S, Reiman D, Clancy E, Zielinski M, Steinegger M, Pacholska M, Berghammer T, Bodenstein S, Silver D, Vinyals O, Senior AW, Kavukcuoglu K, Kohli P, Hassabis D. 2021. Highly accurate protein structure prediction with AlphaFold. Nature 596, 583–589.

Kachanovsky DE, Filler S, Isaacson T, Hirschberg J . 2012. Epistasis in tomato color mutations involves regulation of phytoene synthase 1 expression by cis-carotenoids. Proceedings of the National Academy of Sciences of the United States of America 109, 19021–19026.

Lau OS, Deng XW . 2012. The photomorphogenic repressors COP1 and DET1: 20 years later. Trends Plant Sci 17, 584–593.

Li H, Durbin R . 2009. Fast and accurate short read alignment with Burrows-Wheeler transform. Bioinformatics 25, 1754–1760.

Li H, Handsaker B, Wysoker A, Fennell T, Ruan J, Homer N, Marth G, Abecasis G, Durbin R . 2009. The Sequence Alignment/Map format and SAMtools. Bioinformatics 25, 2078–2079.

Lin Z, Akin H, Rao R, Hie B, Zhu Z, Lu W, Smetanin N, Verkuil R, Kabeli O, Shmueli Y, dos Santos Costa A, Fazel-Zarandi M, Sercu T, Candido S, Rives A . 2023. Evolutionary-scale prediction of atomic-level protein structure with a language model. Science 379, 1123–1130.

Lindgreen S. 2012. AdapterRemoval: easy cleaning of next-generation sequencing reads. BMC Res Notes 5, 337.

Llorente B, Martinez-Garcia JF, Stange C, Rodriguez-Concepcion M . 2017. Illuminating colors: regulation of carotenoid biosynthesis and accumulation by light. Curr Opin Plant Biol 37, 49–55.

Maass D, Arango J, Wust F, Beyer P, Welsch R . 2009. Carotenoid crystal formation in Arabidopsis and carrot roots caused by increased phytoene synthase protein levels. PLOS ONE 4, e6373.

Mariani V, Kiefer F, Schmidt T, Haas J, Schwede T . 2011. Assessment of template based protein structure predictions in CASP9. Proteins 79 Suppl 10, 37–58.

Mathur J, Koncz C . 1998. Callus culture and regeneration. In: Martinez-Zapater J, Salinas J, eds. Arabidopsis protocols: Methods in molecular biology, Vol. 82. Totowa: Humana Press, Inc, 31-34.

McQuinn RP, Leroux J, Sierra J, Escobar-Tovar L, Frusciante S, Finnegan EJ, Diretto G, Giuliano G, Giovannoni JJ, Leon P, Pogson BJ . 2023. Deregulation of zeta-carotene desaturase in Arabidopsis and tomato exposes a unique carotenoid-derived redundant regulation of floral meristem identity and function. Plant J 114, 783–804.

Mechin V, Damerval C, Zivy M. 2007. Total protein extraction with TCA-acetone. Methods Mol Biol 355, 1–8.

Mirdita M, Schutze K, Moriwaki Y, Heo L, Ovchinnikov S, Steinegger M . 2022. ColabFold: making protein folding accessible to all. Nat Methods 19, 679–682.

Moreno JC, Mi J, Alagoz Y, Al-Babili S . 2021. Plant apocarotenoids: from retrograde signaling to interspecific communication. Plant J 105, 351–375.

Nayak JJ, Anwar S, Krishna P, Chen ZH, Plett JM, Foo E, Cazzonelli CI . 2022. Tangerine tomato roots show increased accumulation of acyclic carotenoids, less abscisic acid, drought sensitivity, and impaired endomycorrhizal colonization. Plant Sci 321, 111308.

Ossowski S, Schneeberger K, Clark RM, Lanz C, Warthmann N, Weigel D . 2008. Sequencing of natural strains of Arabidopsis thaliana with short reads. Genome Research 18, 2024–2033.

Osterlund MT, Hardtke CS, Wei N, Deng XW . 2000. Targeted destabilization of HY5 during light-regulated development of Arabidopsis. Nature 405, 462–466.

Pandit J, Danley DE, Schulte GK, Mazzalupo S, Pauly TA, Hayward CM, Hamanaka ES, Thompson JF, Harwood HJ, Jr. 2000. Crystal structure of human squalene synthase. A key enzyme in cholesterol biosynthesis. J Biol Chem 275, 30610–30617.

Park H, Kreunen SS, Cuttriss AJ, DellaPenna D, Pogson BJ . 2002. Identification of the carotenoid isomerase provides insight into carotenoid biosynthesis, prolamellar body formation, and photomorphogenesis. Plant Cell 14, 321–332.

Pfaffl MW . 2001. A new mathematical model for relative quantification in real-time RT-PCR. Nucleic Acids Res 29, e45.

Porra RJ, Thompson WA, Kriedemann PE . 1989. Determination of accurate extinction coefficients and simultaneous equations for assaying chlorophylls a and b extracted with four different solvents: verification of the concentration of chlorophyll standards by atomic absorption spectroscopy. Biochim. Biophys. Acta 975, 384–394.

Rodriguez-Villalon A, Gas E, Rodriguez-Concepcion M . 2009. Colors in the dark: a model for the regulation of carotenoid biosynthesis in etioplasts. Plant Signal Behav 4, 965–967.

Ruiz-Sola MA, Coman D, Beck G, Barja MV, Colinas M, Graf A, Welsch R, Rutimann P, Buhlmann P, Bigler L, Gruissem W, Rodriguez-Concepcion M, Vranova E . 2016. Arabidopsis GERANYLGERANYL DIPHOSPHATE SYNTHASE 11 is a hub isozyme required for the production of most photosynthesis-related isoprenoids. New Phytol 209, 252–264.

Schaub P, Rodriguez-Franco M, Cazzonelli CI, Álvarez D, Wüst F, Welsch R. 2018. Establishment of an Arabidopsis callus system to study the interrelations of biosynthesis, degradation and accumulation of carotenoids. PLOS ONE 13, e0192158.

Schneeberger K, Ossowski S, Lanz C, Juul T, Petersen AH, Nielsen KL, Jorgensen JE, Weigel D, Andersen SU . 2009. SHOREmap: simultaneous mapping and mutation identification by deep sequencing. Nat Methods 6, 550–551.

Schrödinger LLC. 2015. The PyMOL Molecular Graphics System.

Shumskaya M, Bradbury LM, Monaco RR, Wurtzel ET . 2012. Plastid localization of the key carotenoid enzyme phytoene synthase is altered by isozyme, allelic variation, and activity. Plant Cell 24, 3725–3741.

Shumskaya M, Wurtzel ET. 2013. The carotenoid biosynthetic pathway: thinking in all dimensions. Plant Sci 208, 58–63.

Stephenson PG, Fankhauser C, Terry MJ . 2009. PIF3 is a repressor of chloroplast development. Proc Natl Acad Sci U S A 106, 7654–7659.

Toledo-Ortiz G, Huq E, Rodriguez-Concepcion M . 2010. Direct regulation of phytoene synthase gene expression and carotenoid biosynthesis by phytochrome-interacting factors. Proc Natl Acad Sci U S A 107, 11626–11631.

Toledo-Ortiz G, Johansson H, Lee KP, Bou-Torrent J, Stewart K, Steel G, Rodríguez-Concepción M, Halliday KJ. 2014. The HY5-PIF Regulatory Module Coordinates Light and Temperature Control of Photosynthetic Gene Transcription. PLOS Genetics 10, e1004416.

Welsch R, Arango J, Bar C, Salazar B, Al-Babili S, Beltran J, Chavarriaga P, Ceballos H, Tohme J, Beyer P . 2010. Provitamin A accumulation in cassava (Manihot esculenta) roots driven by a single nucleotide polymorphism in a phytoene synthase gene. Plant Cell 22, 3348–3356.

Welsch R, Beyer P, Hugueney P, Kleinig H, von Lintig J . 2000. Regulation and activation of phytoene synthase, a key enzyme in carotenoid biosynthesis, during photomorphogenesis. Planta 211, 846–854.

Welsch R, Wust F, Bar C, Al-Babili S, Beyer P. 2008. A third phytoene synthase is devoted to abiotic stress-induced abscisic acid formation in rice and defines functional diversification of phytoene synthase genes. Plant Physiology 147, 367–380.

Welsch R, Zhou X, Yuan H, Álvarez D, Sun T, Schlossarek D, Yang Y, Shen G, Zhang H, Rodriguez-Concepcion M, Thannhauser TW, Li L. 2018. Clp Protease and OR Directly Control the Proteostasis of Phytoene Synthase, the Crucial Enzyme for Carotenoid Biosynthesis in Arabidopsis. Molecular Plant 11, 149–162.

Xu X, Chi W, Sun X, Feng P, Guo H, Li J, Lin R, Lu C, Wang H, Leister D, Zhang L 2016. Convergence of light and chloroplast signals for de-etiolation through ABI4-HY5 and COP1. Nat Plants 2, 16066.

Zhou X, Rao S, Wrightstone E, Sun T, Lui ACW, Welsch R, Li L . 2022. Phytoene Synthase: The Key Rate-Limiting Enzyme of Carotenoid Biosynthesis in Plants. Front Plant Sci 13, 884720.

Zhou X, Welsch R, Yang Y, Alvarez D, Riediger M, Yuan H, Fish T, Liu J, Thannhauser TW, Li .L 2015. Arabidopsis OR proteins are the major posttranscriptional regulators of phytoene synthase in controlling carotenoid biosynthesis. Proc Natl Acad Sci U S A 112, 3558–3563.

