## Supplementary Figures and Tables for "Reducing PSY activity fine tunes threshold levels of a *cis*-carotene-derived signal that regulates the PIF3/HY5 module and plastid biogenesis"

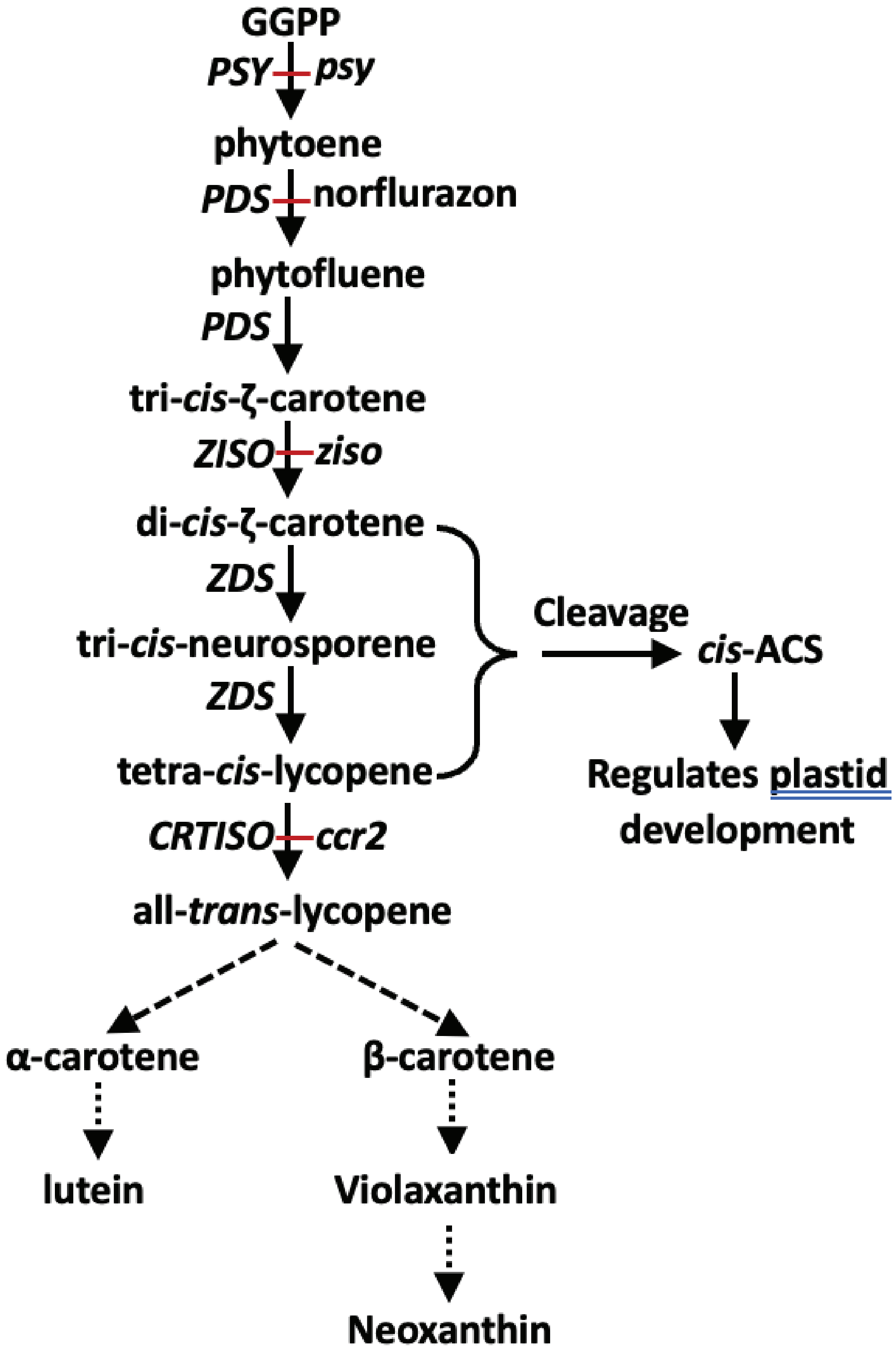


**Supplemental Fig. 1: Simplified diagram of the carotenoid biosynthesis pathway.** Genes encoding the catalysing enzymes of each step before all-trans-lycopene are labelled to the left of the pathway and mutants are labelled to the right (red line indicates blocked step). GGPP: geranylgeranyl diphosphate; PSY: phytoene synthase; phytoene: cis-phytoene; phytofluene: di-cis-phytofluene; PDS: phytoene desaturase; ZISO: 15-cis-ζ-carotene isomerase; ZDS: ζ-carotene desaturase; CRTISO: carotenoid isomerase. Norflurazon is an herbicide that inhibits PDS activity. cis-ACS: cis-carotene derived apocarotenoid signal (ACS).

**A**

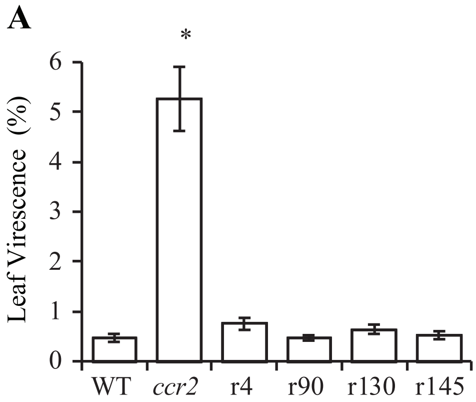


**Supplemental Fig. 2: Percentage of leaf virescence leaves in rccr2 lines reverted the leaf-yellowing phenotype in ccr2 and led to albino phenotype in one line.** A) Percentage of yellow leaf area of WT, ccr2 and rccr2 (r) lines. Plants were grown under 8-h photoperiod. Star denotes significant difference in comparison to WT ($p < 0.05$ in one-way ANOVA).

**A**

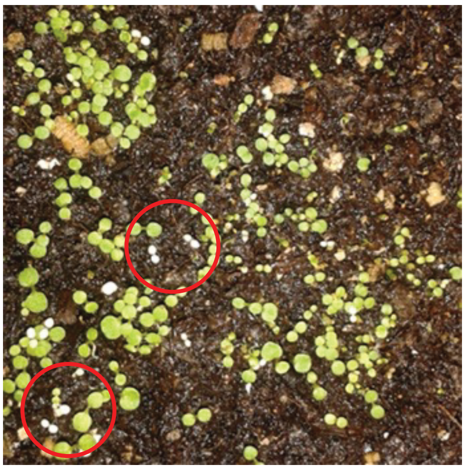

**B**

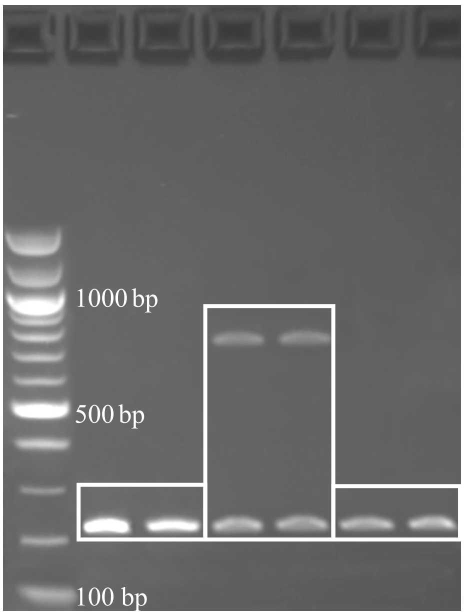


**C**

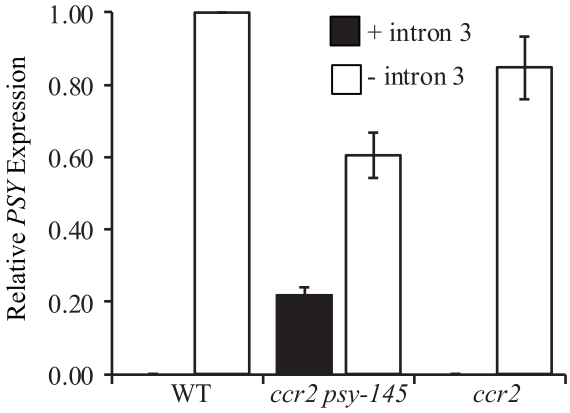


**Supplemental Fig. 3: Characterisation of ccr2 psy-145 and alternative splicing of PSY.** A) Albino seedlings displayed by rccr2–145 (ccr2 psy-145 - red circle) seedlings were grown under a 16-h photoperiod. B) Reverse transcription PCR (RT-PCR) showing alternative splicing of intron 3 in ccr2 psy–145 leaf tissues. Exons 2 and 3 were partially amplified, and products were visualised on a 1% agarose gel. Spliced shows 221 bp (WT and ccr2) and unspliced 762 bp (ccr2 psy–145) amplicons. 100 bp ladder is displayed. C) Quantitative RT-PCR (qRT-PCR) of PSY mRNA amplifying part of intron 3 (162 bp) in ccr2 psy–145 (+) or a region (155bp) spanning exon 2 and 3 when intron 3 is spliced out in WT and ccr2 leaf tissues. Standard error bars are shown (n=10). RNA levels were normalized to Protein Phosphatase 2A (AT1G13320) reference gene validated for mRNA normalisation in leaves using Cyclophilin (At2g29960) and TIP41 (At4g34270) secondary reference genes (Cazzonelli et al., 2014; Cazzonelli et al., 2010b).

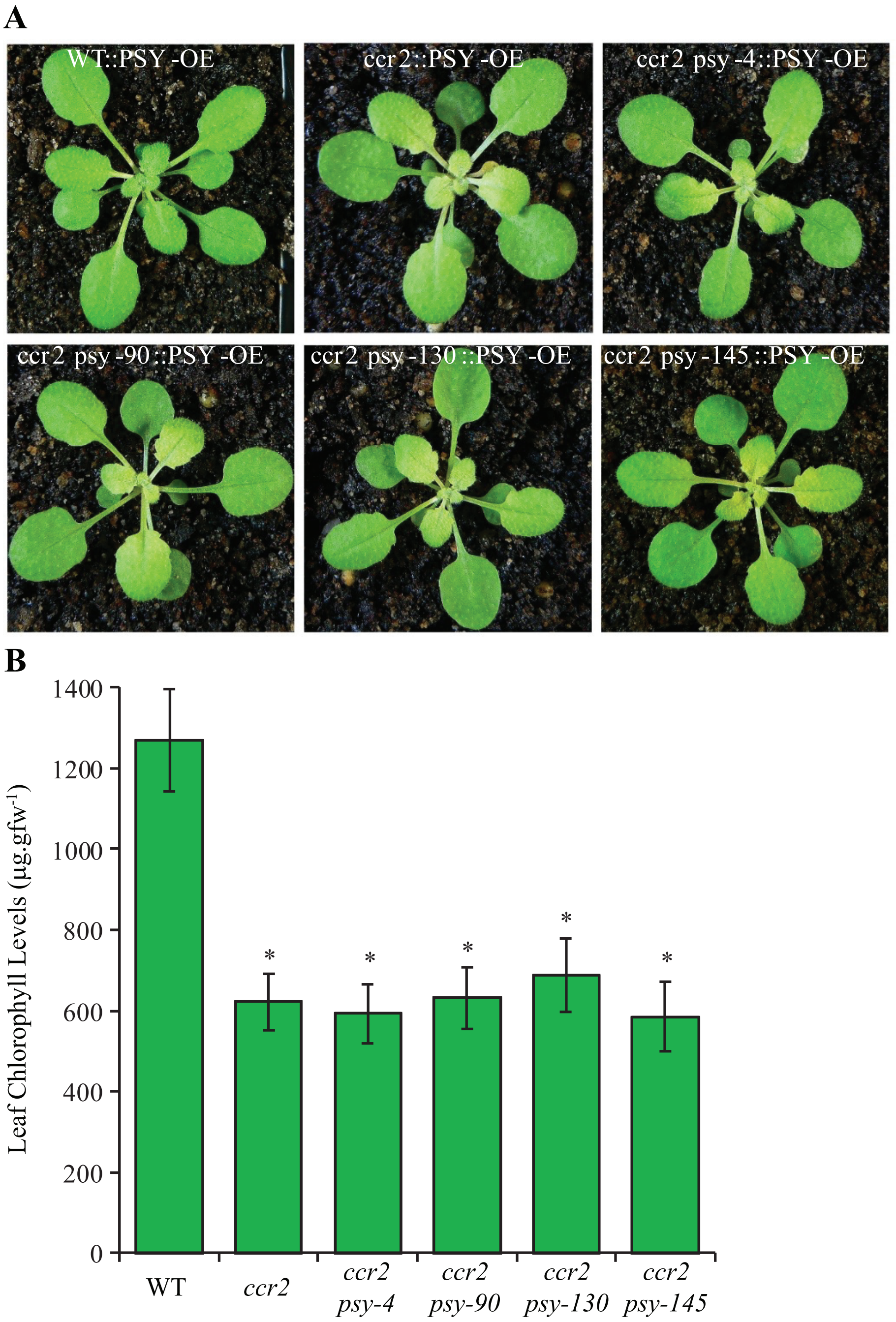


**Supplemental Fig. 4: Overexpression of functional PSY restores ccr2 mutant phenotypes in ccr2 psy double mutant lines.** A) Representative images of rosettes show yellow virescence in newly emerged leaves from ccr2, and ccr2 psy lines compared to the WT control and WT::PSY-OE. T4 generation transgenic plants plants were grown under a 10-h photoperiod for 3 weeks and images from 50-100 plants analysed for five independent lines. PSY-OE; PSY overexpression. B) Total chlorophyll in leaves from T4 generation transgenic plants. Plants were grown under an 8-h photoperiod. Error bars denote standard error of means (n=5). Star denotes significant difference in comparison to WT-OE ($p < 0.05$ in one-way ANOVA). YL; yellow leaf.

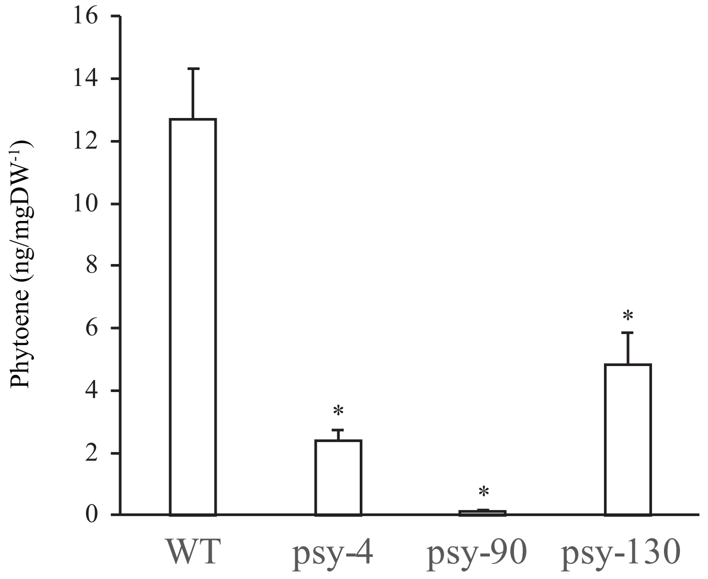


**Supplemental Fig. 5: Split ubiquitin assays showing phytoene levels generated in yeast cells expressing PSY+AtGGPPS11 combinations.** Phytoene levels were measured using HPLC. Error bars indicate standard error of means (n=3). Star denotes a significant difference in comparison to WT ($p < 0.05$; one-way ANOVA).

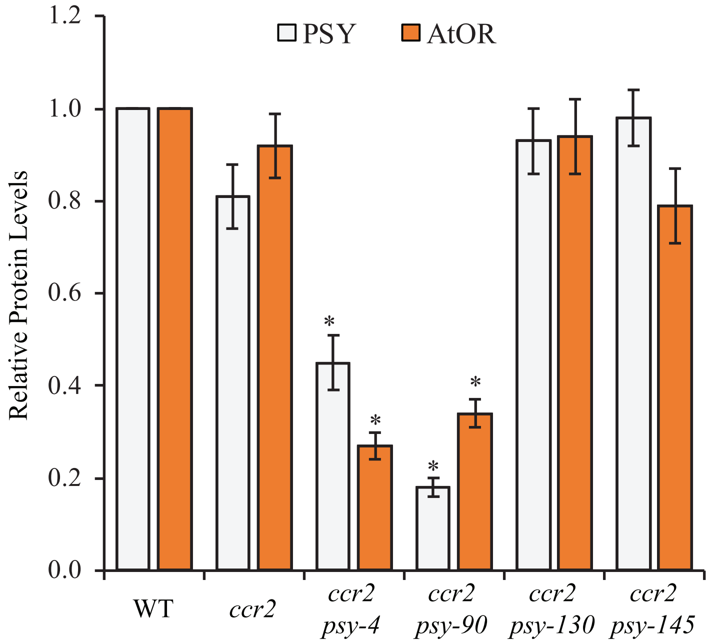


**Supplemental Fig. 6: ELISA of PSY and AtOR protein levels.** Each well was coated with total protein from Arabidopsis leaf tissue and incubated with anti-PSY or anti-OR antibodies. Following the addition of substrate solution containing O-phenylenediamine dihydrochloride and H2O2 absorbances were measured at 492 nm. Values were averaged from five biological replicates and Error bars denote standard error of means (n=5). Stars denote the significant difference in comparison to WT ($p < 0.05$; one-way ANOVA).

### Supplemental Material

#### Supplemental Table 1: Primers used in this study

| Name | Sequence | Accession |
|---|---|---|
| **Sequencing** | | |
| PSY4F<br>PSY4R | CTCTGTCTTATCTTACTTCC<br>TAAACCCTTCCTCTTCTC | At5g17230 |
| PSY90F<br>PSY90R | ACCGGTTTCTTGATTCACA<br>TTCCACTTTCCTCTCGCT | At5g17230 |
| PSY130F<br>PSY130R | GAACACCAAGCATCCAAA<br>GTCTCGAAATGGCTGCAA | At5g17230 |
| PSY145F<br>PSY145R | GGTCTTCTTCTTATGACC<br>CATCACTTTATCCTACAA | At5g17230 |
| **Cloning** | | |
| PSY145FC<br>PSY145RC | CTTTGCTTATGACACCCG<br>CAGAATATCGACCGGGTATC | At5g17230 |
| **qRT-PCR** | | |
| PSY145eFq<br>PSY145eRq | GGCAATCTACGGTAAGTTAC<br>GCAACTGTATCAGCGAGA | At5g17230 |
| PSY145iFq<br>PSY145iRq | CCAATGGTTGAAGAGCTG<br>CAGCTTCAACTTCTCTTG | At5g17230 |
| PIF3F<br>PIF3R | TTGGCTCGGGTAATAGTCTCGATG<br>CCTGCTTCCTTTCTTCCATCTCCT | At1g09530 |
| HY5F<br>HY5R | GAGAAAGAGAACAAGCGGCTGAAG<br>AGCATCTGGTTCTCGTTCTGAAGA | At5g11260 |
| PP2A-cc1F<br>PP2A-cc1R | CTTCGTGCAGTATCGCTTCTC<br>ATTGGAGAGCTTGATTTGCG | At1g13320 |
| **Yeast split ubiquitin system** | | |
| B1-PSY<br>B2-PSY | B1-TCTTTTGTAAGGAACCGAAGTAG<br>B2-TATCGATAGTCTTGAACTTGAAG | At5g17230 |
| B1-AtOR<br>B2-AtOR | B1-GCCGATAAATTCGCTTCCGGG<br>B2-ATCGAAAGGGTCGATACGAGGATC | At5g61670 |
| B1-AtGGPPS11<br>B2-AtGGPPS11 | B1-TCTTCTTCCGTTGTTACAAAAG<br>B2-GTTCTGTCTATAGGCAATGTAATT | At4g36810 |
| B1 linker<br>B2 linker | ACAAGTTTGTACAAAAAAGCAGGCTCTCCAACCACCATG<br>TCCGCCACCACCAACCACTTTGTACAAGAAAGCTGGGTA | No Applicable |

**Supplemental Table 2: Genomic information of the mutations identified in *PSY* from *rccr2* lines.** SNP; single nucleotide polymorphism.

| ***rccr2* lines** | **SNP position** | **Base change** | **Gene Location** | **Exon Position** | **Amino acid change** |
|---|---|---|---|---|---|
| *rccr2*$^{-4}$ | 5660555 | C→T | CDS | 4 | M→I |
| *rccr2*$^{-90}$ | 5660151 | C→T | CDS | 5 | A→T |
| *rccr2*$^{-130}$ | 5660938 | G→A | CDS | 4 | P→S |
| *rccr2*$^{-145}$ | 5661514 | C→T | Splice site | Exon2/intron splice site | Truncated protein |
